# Structural basis of an EMC:Spf1 insertase-dislocase complex in the eukaryotic endoplasmic reticulum

**DOI:** 10.1101/2025.08.07.669073

**Authors:** Carolin J Klose, Jesuraj R Prabu, Emma J Fenech, Bastian Bräuning, Ida Baydar, Barbara Steigenberger, Susanne von Gronau, Sivan Arad, Christine R Langlois, Maya Schuldiner, Brenda A Schulman, Matthias J Feige

## Abstract

Most eukaryotic membrane proteins are inserted into the membrane at the endoplasmic reticulum (ER). This essential but error-prone process relies on molecular quality control machineries to prevent mistargeting and incorrect structure formation. Here we show that the ER membrane protein complex (EMC) forms an evolutionarily conserved supercomplex with the P5A-ATPase Spf1/ATP13A1. This supercomplex combines the transmembrane domain (TMD) insertase function of the EMC and the TMD dislocase activity of Spf1 in a single entity. Our cryo-EM structure of this supercomplex shows that EMC and Spf1 form a shared intramembrane cavity for substrate engagement and reveals that the ATPase cycle of Spf1 regulates access to this cavity. Together, our study suggests that proteins with opposing biochemical activities in membrane protein biogenesis – insertion versus dislocation – form an integrated molecular machine in eukaryotes to proofread membrane protein insertion and topogenesis.

## Introduction

Integral membrane proteins account for about one quarter of all eukaryotic genes^2,3^. They mediate essential processes in cells, including signal transduction, organelle tethering, lipid homeostasis, and the transport of ions, nutrients and entire proteins. The biogenesis of membrane proteins is a complex process that relies on correct protein targeting, insertion into the lipid bilayer and structure formation within and outside the membrane. Eukaryotic cells thus have evolved multiple, complementary systems to safeguard targeting and insertion of membrane proteins into the lipid bilayer. For the secretory pathway, which acts as a hub for lipid bilayer integration and structure formation of most membrane proteins, targeting mechanisms include the signal recognition particle (SRP) pathway^2,4^, the guided entry of tail-anchored (TA) proteins (GET) system^5^ and the SRP-independent targeting (SND) pathway^6^. Downstream of the SRP-pathway, the Sec61 translocon was initially identified as the principal conduit for co-translational insertion of nascent polypeptides into the into the Endoplasmic Reticulum (ER) membrane^7-9^. However, beyond Sec61, a growing number of specialized machineries have been discovered that mediate the insertion and folding of structurally diverse membrane protein clients. Recent findings show that these machineries can assemble into large multi-subunit complexes, such as the multipass-translocon^10^ and the holo-insertase complex^11^, to accommodate diverse clients.

A key machinery for membrane protein biogenesis is the ER membrane protein complex (EMC), which has emerged as a central player in membrane protein biogenesis and the maintenance of cellular homeostasis^12-15^. The EMC is an evolutionarily conserved complex in eukaryotes, comprising eight subunits in yeast and nine in humans^16^. Central to the EMC’s function is its ability to insert TMDs into the ER membrane, particularly terminal ones and those with moderate hydrophobicity^14,15^. Beyond its insertase function, the EMC has been shown to act as a chaperone for membrane proteins^17-19^ and serves as a central hub in membrane protein biogenesis and quality control by interacting with multiple further quality control factors^11,19,20^.

While organellar membrane protein insertases generally do display preferences for certain clients, they often allow for promiscuous insertions to accommodate a spectrum of client helix properties. As a consequence, they are error-prone and can mistarget and misinsert clients into the wrong organelle. This explains the emergence of quality control machineries downstream of membrane protein insertion. In mitochondria, for example, mistargeted TA proteins are extracted from the outer membrane by the AAA-ATPase Msp1 to be degraded at the ER^21,22^. Similarly, the EMC may erroneously integrate mitochondrial TA proteins into the ER membrane^1,23-25^. Here, the P5A-ATPase Spf1 (yeast)/ ATP13A1 (human) functions as a dislocase, surveilling the integration of TA proteins, as well as erroneously integrated signal sequences and TMDs^26^. This extraction activity maintains organellar integrity and can give transmembrane and signal sequence containing proteins another chance to obtain their destined localization and native structure^27,28^. The dislocase function of Spf1/ATP13A1 is driven by its ATPase cycle, referred to as the Post-Albers cycle^29,30^. While most P-type ATPases transport cargo from the cytosol into the ER lumen or extracellular space, Spf1 operates in the opposite direction, extracting its polypeptide cargo from the ER membrane to the cytosol^26,31^. Thus, by acting together, insertases, chaperones, and dislocases support the fidelity of organelle-specific membrane protein biogenesis. The mechanistic details of this interplay, however, remain largely uncharacterized.

Given EMC’s central role in this network, we aimed to identify regulators and interactors that contribute to this key functional interplay. Our data reveal that the transmembrane helix dislocase Spf1/ATP13A1 is a stoichiometric interactor of the EMC in both yeast and human cells at endogenous levels. Cryo-electron microscopy (cryo-EM) structures of the EMC:Spf1 complex reveal a close juxtaposition of the EMC insertase and Spf1 dislocase cavities and show how the ATPase activity of Spf1 modulates the conformation and architecture of the overall complex. Our study reveals how molecular machines with opposing functions: protein insertion and extraction, can act as a single integrated assembly with dual functionality, suggesting a new paradigm in membrane protein biogenesis.

## Results

### The EMC and Spf1 form a stoichiometric complex in cells

The multi-subunit architecture of the EMC and recent studies on this complex^11,17,19,20^ indicate that it is a central hub for membrane protein biogenesis rather than a stand-alone molecular machine. To gain a comprehensive view of the endogenous cofactors and modulators that associate with the EMC, we tagged one of its core subunits (Emc5) in *Saccharomyces cerevisiae* (from here on called yeast) with a C-terminal triple FLAG-tag and performed immunoprecipitation coupled to mass spectrometry (IP-MS) to define the EMC interactome. We employed a previously established mild isolation strategy^16^, using the non-ionic detergent digitonin to also preserve weak interactions (Fig. 1a, Extended Data Fig. 1a). This approach identified approximately 250 interactors of the yeast EMC, including previously characterized ones like the chaperone Ilm1^16^, as well as proteins involved in ER targeting and insertion, the OST (oligosaccharyl transferase) complex, ERAD (ER associated degradation)-related proteins, ER-shaping proteins, and numerous factors involved in lipid and sterol metabolism (Fig. 1b, Supplementary Table 1, SI Fig. 1). Among these, Spf1 stood out with an enrichment level comparable to the EMC subunits themselves (Fig. 1b), in agreement with a previous report of this interaction in an interactomics experiment^16^.

**Figure 1.**
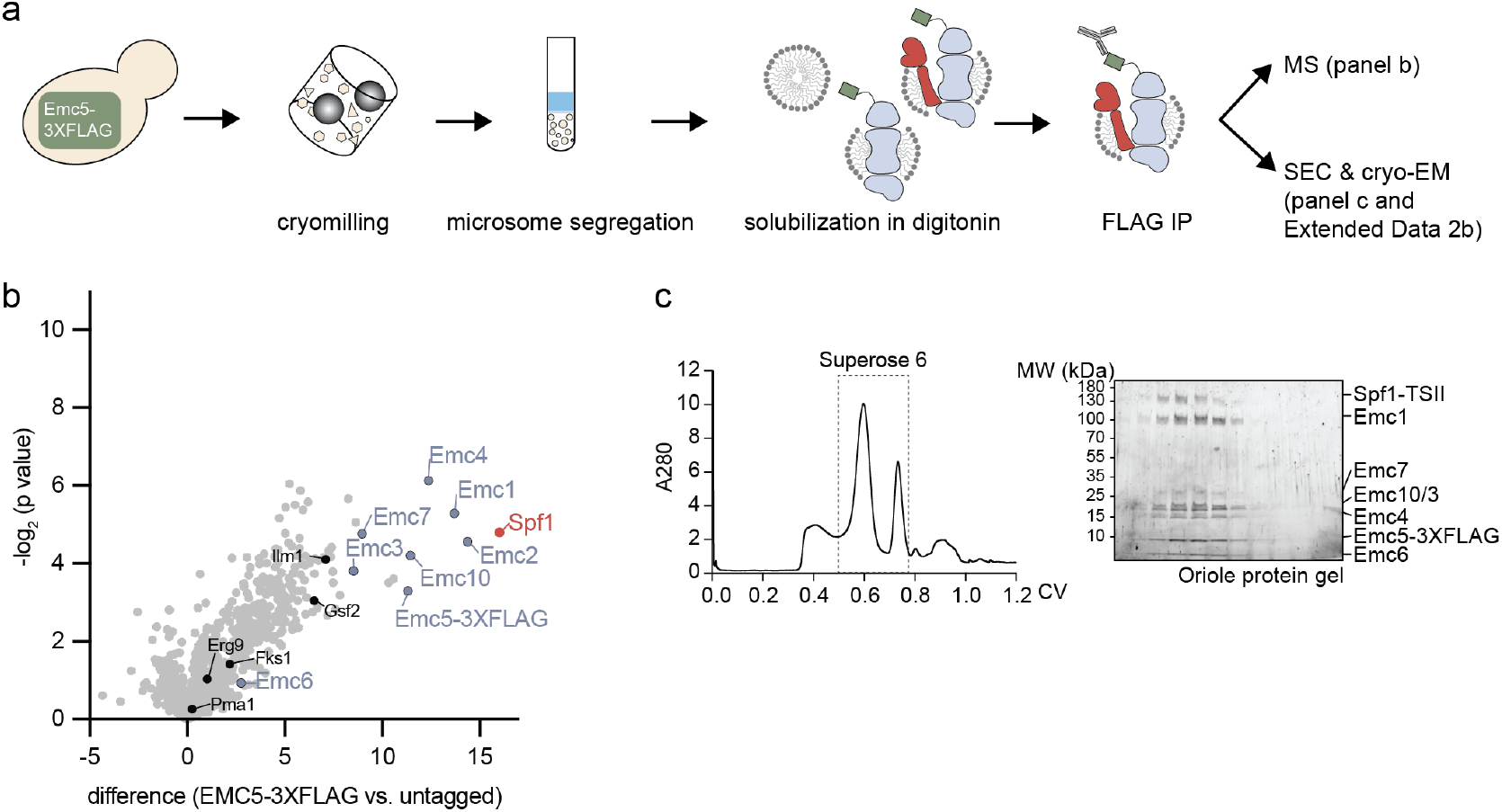
EMC and Spf1 form a stoichiometric complex. **(A)** Procedure for mild isolation of endogenously tagged Emc5-3XFLAG from *Saccharomyces cerevisiae* by cryogenic disruption of the cell wall, centrifugal isolation of microsomes and solubilization in digitonin before immunocapture of the 3XFLAG tag. **(B)** The interactome of Emc5-3XFLAG reveals Spf1 (red) as the most prevalent interaction partner of the EMC in yeast. EMC subunits are labeled in light blue and previously identified interactors in black^16^. **(C)** In a large-scale purification of the EMC, Spf1 co-elutes in a stoichiometric manner after a single pulldown on Emc5 and size exclusion chromatography. The figure shows the UV 280 nm absorption trace of the SEC and SDS-PAGE analysis of indicated fractions stained for whole protein. The full purification is shown in Extended Data Fig. 1d.

This pronounced interaction might be unexpected, since the function of Spf1 as a transmembrane helix dislocase in the ER membrane^26^ is biochemically opposing to the role of the EMC as an insertase. Of note, similar phenotypes after knockout of either have been described, related to defects in sterol metabolism^32-34^. Furthermore, recent studies discussed an overlap in client spectra between the human orthologs hEMC and ATP13A1^25,35,36^ which may indicate synergies. Thus, to validate this interaction, we endogenously tagged Spf1 with a TwinStrepII (TSII)-tag and confirmed its presence in an anti-Emc5-FLAG IP by western blotting (Extended Data Fig. 1b). Additionally, we performed the reciprocal IP-MS experiment on endogenously tagged Spf1 and found the EMC subunits to be amongst the most highly enriched interactors (Extended Data Fig. 1c, Supplementary Table 1, SI Fig. 1).

To further characterize the EMC:Spf1 complex, we purified the endogenous EMC from yeast in digitonin at a large scale. Following a pulldown of Emc5 and subsequent size exclusion chromatography (SEC), Spf1 was present at levels comparable to the EMC subunits (Fig. 1c, Extended Data Fig. 1d, e), providing direct evidence for stable endogenous complex formation between the EMC and Spf1.

### Cryo-EM structure of the EMC:Spf1 complex reveals juxtaposed functional sites

To gain insights into the molecular details of the EMC:Spf1 complex, we obtained large homogeneous amounts of the yeast complex by recombinant production in insect cells and a tandem affinity purification followed by SEC (Fig. 2a, Extended Data Fig. 2a). This allowed us to determine the structure of the complex by cryo-EM to a global resolution of 3.8 Å for the EMC and 4.2 Å for Spf1 (Fig. 2b, SI Fig. 2). As a validation, we additionally purified the endogenous complex from yeast via endogenously tagged Emc5 and obtained a low-resolution cryo-EM map (global resolution 8.7 Å) that was fully consistent with the recombinant complex (Extended Data Fig. 3a-d, SI Fig. 3).

**Figure 2.**
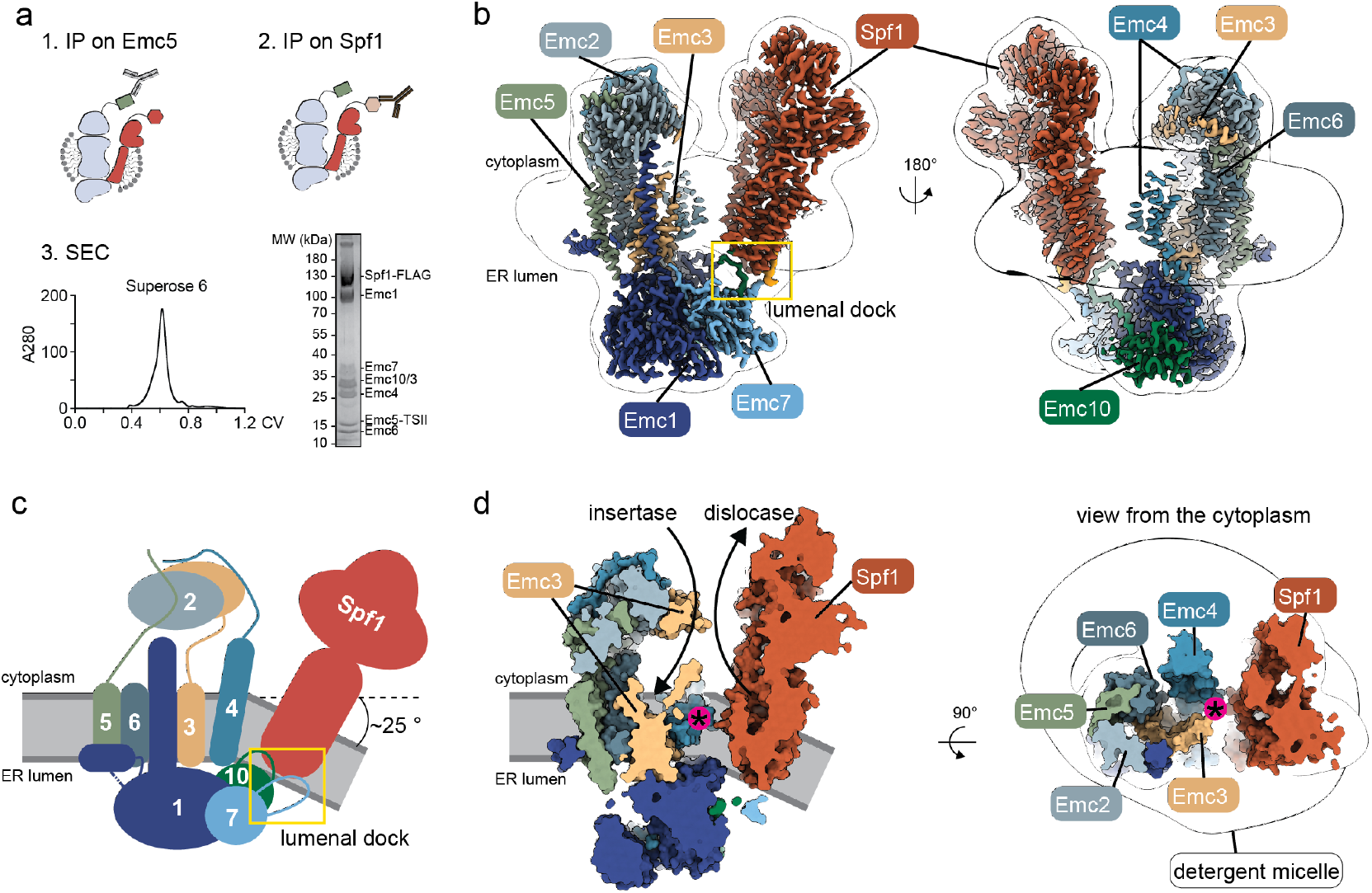
The cryo-EM structure of the EMC:Spf1 complex reveals a large composite intramembrane cavity encompassing both client helix engagement sites. **(A)** Recombinant expression and purification of the EMC:Spf1 complex in insect cells via a tandem affinity pulldown and SEC yielding a homogeneous sample for cryo-EM as shown on the Coomassie stained SDS-PAGE gel. **(B)** Cryo-EM map of the EMC:Spf1 complex colored according to subunit identity. The out-line of a low-pass filtered map is overlaid to show the borders of the detergent micelle. The main interaction interface between the EMC and Spf1, the lumenal dock, is marked with a yellow box. **(C)** Model cartoon depicting the subunit architecture of the EMC:Spf1 complex and high-lighting the bend of approximately 25° in the detergent micelle and the lumenal dock. **(D)** The insertase site of the EMC is facing the dislocase site of Spf1 inside the shared in-tramembrane cavity. View of the cavity from the membrane (left) and top view from the cytoplasm (right) show the size and continuity of the composite cavity which is marked with a magenta asterisk.

In the EMC:Spf1 complex, all EMC subunits (Emc1-7, Emc10) were present in an arrangement similar to previously reported structures of the yeast EMC (Extended Data Fig. 2b, RMSD to 7KRA: 0.9 Å, to 7KTX: 1.1 Å, to 6WB9: 1.2 Å)^37,38^. Extending this previous work, we resolved additional C-terminal residues for the subunits Emc4, Emc7 (also named Sop4) and Emc10^37,38^. Notably, compared to the prior structures of isolated EMC complexes, the cytoplasmic cap, composed of Emc2, 3 and 5 and known to be involved in initial substrate engagement and selection^1,39^, was tilted and displaced by approximately 5 Å (Extended Data Fig. 2c). This displacement was most pronounced in the cytoplasmic helices of Emc3, the subunit which is homologous to the evolutionarily conserved Oxa1/Alb3/YidC family of insertases^40^.

The overall architecture of Spf1 in the complex was also consistent with previously reported Spf1 structures (Extended Data Fig. 4a, RMSD to 6XMP: 1.0 Å)^26,31^. Spf1 is a P-type ATPase whose ability to translocate substrates is coupled to ATP hydrolysis and cycling between two major conformational states: E1 and E2 ^26,31^. In the absence of any nucleotide-state stabilizing analog, Spf1 in complex with the EMC is in an E1 conformation with its intramembrane cleft facing the cytosol. We observed the most prominent interaction surface between the ER lumenal parts of the proteins, involving Emc7, 10 and the lumenal extensions of Spf1. This “lumenal dock” buries around 413.7 Å^2^ of surface area (356.2 Å^2^ with Emc10, 57.5 Å^2^ with Emc7) (Fig. 2b, c). Formation of this interface requires the EMC and Spf1 to approach each other at an angle, inducing a bend in the detergent micelle of around 25°, which was also visible in our endogenous map (Fig. 2b, c, Extended Data Fig. 3b).

A well-studied function of the EMC is the insertion of terminal transmembrane helices into the lipid bilayer^14,15,41^ and it is also implicated in the insertion of TMDs separated by only short hydrophilic loops^42^. These functions have largely been attributed to an intramembrane hydrophilic vestibule enclosed by the subunits Emc3, 4 and 6, often referred to as the “gated cavity”^37,38,43,44^. Spf1 has been shown to bind substrate helices within its intramembrane cleft^26,31^. In the EMC:Spf1 complex structure, these two validated functional sites for substrate engagement directly face each other, forming a composite cavity inside the membrane (Fig. 2d, Extended Data Fig. 4c, d). The cavity resembles a funnel shape spanning the whole transmembrane region and has a width of 25 Å at the ER lumenal and 50 Å at the cytoplasmic side of the membrane (Extended Data Fig. 4d). The cavity is laterally lined by the Emc4 gate on one side, which approaches Spf1 at a 5 Å distance on the lumenal side (buried surface area of 52.1 Å^2^). On the opposing side, Emc7 delimits the central vestibule, for which we resolved 12 more residues than previously reported^37,38^, yet could not detect the transmembrane domain and following C-terminal residues. Together, our data reveal a large composite intramembrane cavity, which encompasses two validated sites for protein translocation and due to its size could accommodate complex clients with multiple TMDs.

### Molecular architecture of the EMC:Spf1 interface

The main site of interaction between the EMC and Spf1 was confined to an interface at the ER lumenal side, composed of the ER lumenal parts of Spf1’s support domain (TMDs 7 to 10), a loop in Emc7 (F114-L125) and a segment of the Emc10 linker region (K118-Y124), which connects the ER lumenal β-sandwich domain to its structurally unresolved TMD (Fig. 3a). Both subunits, Emc7 and Emc10, are considered accessory rather than core subunits of the EMC, meaning that loss of either leads to mild phenotypes and does not compromise the overall complex stability^17,32^. In fact, Emc7 was first described as a client specific chaperone rather than an EMC subunit^17^ and the function of Emc10 has remained elusive, yet its relevance in human disease points at an important physiological function^45-50^. Additionally, we resolved a digitonin molecule at the lumenal dock, with its sterol backbone lodged into a lateral hydrophobic pocket on Spf1 and its glycosidic chains in close proximity to the Emc7 loop (Fig. 3b). Previous studies have resolved n-dodecyl-β-Dmaltoside (DDM) detergent molecules at the same position on Spf1^26^.

**Figure 3.**
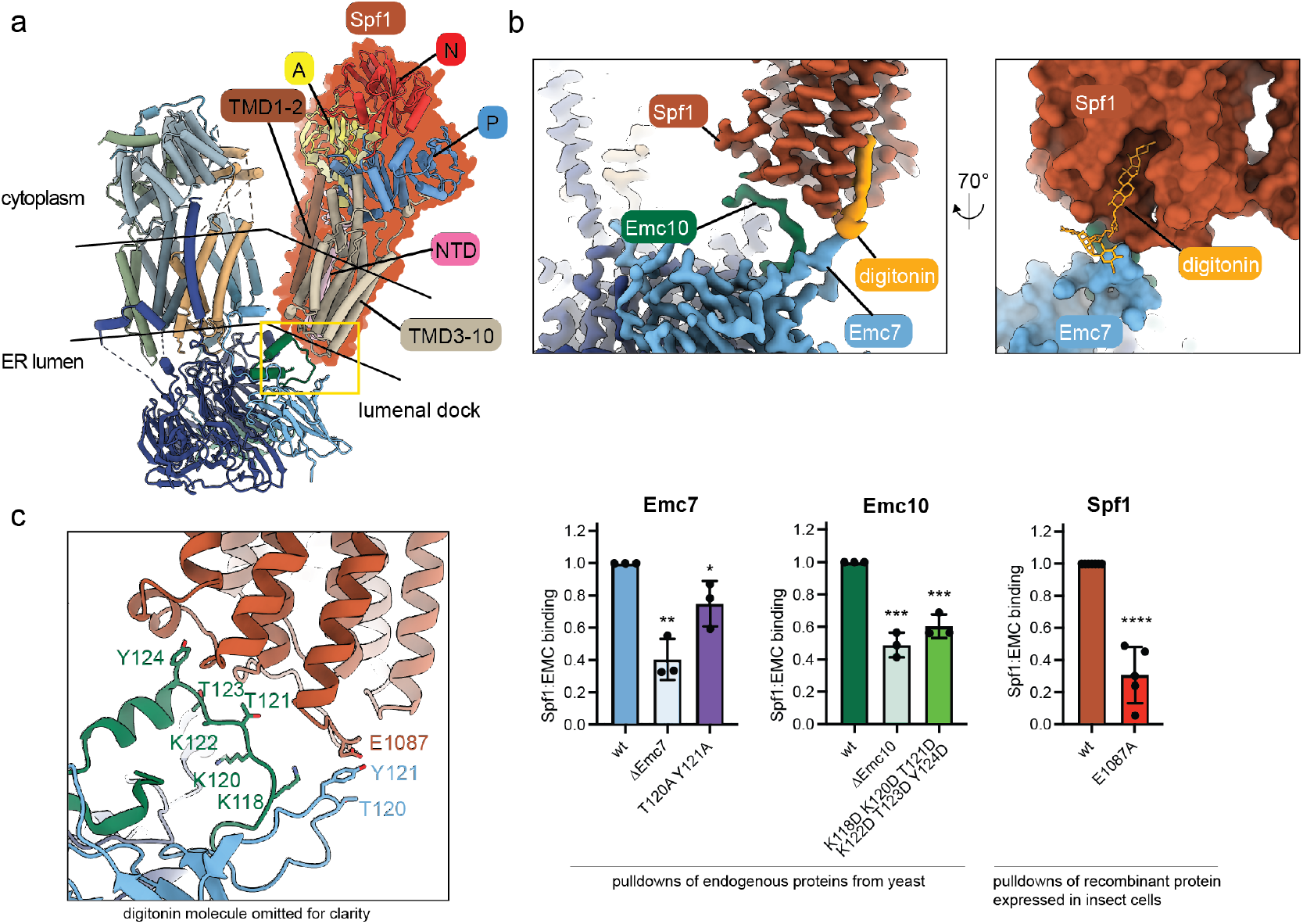
The interaction between the EMC and Spf1 is mediated by an ER-lumenal interface. **(A)** The EMC:Spf1 complex colored according to Spf1 domain architecture with the actuator (A) domain in yellow, the nucleotide binding (N) domain in red, the phosphorylation (P) domain in blue, the N-terminal domain (NTD) in pink and the membrane domain in two shades of brown (see also Extended Data Fig. 4b). The substrate binding site lies between TMD1-2 (brown) and TMD3-10 (sand). The lumenal dock region is marked with a yellow box. **(B)** Left: cryo-EM density of the lumenal dock colored according to protein density. A digitonin molecule (orange) is bound to the back of Spf1, adjacent to its interface with Emc7. Right: detailed view on the bound detergent molecule lodged into a lateral pocket of Spf1. **(C)** Mutational dissection of the lumenal dock. Deletion of Emc7 or Emc10 as well as mutations at the contact sites with Spf1 disrupt the complex in an endogenous setting in yeast. Similarly, introducing a single point mutation in Spf1 strongly destabilizes the EMC:Spf1 complex when co-expressed in insect cells. The digitonin molecule was omitted in the structural representation for clarity. Binding was assessed in three independent replicates (five for Spf1 E1087A) and normalized to the respective wt. (mean ± standard deviation, unpaired Student’s t-test * p < 0.05, ** p < 0.01, *** p < 0.005, **** p < 0.0001).

To probe the key EMC:Spf1 interactions in cells, we generated yeast strains in which one of the core subunits (Emc5) was C-terminally tagged with GFP, Spf1 with a C-terminal triple HA-tag and variants of either Emc7 or Emc10 carried an N-terminal myc-tag (Extended Data Fig. 5a). All genes were manipulated at their endogenous loci under their respective promotor to assess their effects in a setting as close to native conditions as possible. We assessed the integrity of the EMC and its interaction with Spf1 by comparing Emc7 or Emc10 to Emc5 and Spf1 to Emc5 ratios, respectively in Emc5-GFP IPs. Deletion of Emc7 or Emc10 significantly reduced the Emc:Spf1 interaction to 40 % and 48 %, respectively (Fig. 3c, Extended Data Fig. 5b, c), which is in agreement with previous observations^17^. Mutation of two polar residues (T120A Y121A) in the Spf1-contacting loop of Emc7 did not affect incorporation of this Emc7 variant into the EMC but it significantly reduced Spf1 binding (Fig. 3c and Extended Data Fig. 5b). We additionally examined the traversing Emc10-linker region (K118-Y124), which contains multiple positively charged residues that could interact with the negatively charged ER lumenal surface of Spf1 (Extended Data Fig. 5c). By replacing the basic and polar residues in the respective Emc10-linker region with the negatively charged aspartates we expected to disrupt the Emc10:Spf1 interface due to charge repulsion, while not affecting the overall EMC assembly, which was both confirmed by our co-IP data (Fig. 3c, Extended Data Fig. 5d).

Endogenous Spf1 was genetically less accessible than the EMC for introducing mutations in yeast, hence we used our recombinant system to test the effects of complementary mutations in Spf1. The glutamate 1087 in Spf1 faces the polar Emc7 loop which suggested that it may be important for the EMC:Spf1 interaction. Indeed, substitution of this glutamate to alanine significantly reduced the EMC:Spf1 interaction to 30 % (Fig. 3c). This strong effect may also be caused by disrupting binding of the closely bound digitonin molecule, whose glycosidic chains are in close proximity (Fig. 3b).

Taken together, our mutational analysis confirms the structurally observed interaction interface of the EMC:Spf1 supercomplex, but also highlights its robust nature.

### The EMC:Spf1 complex is conserved in human cells

Our data revealed that EMC and Spf1 form a stable complex in yeast, suggesting a potential key functional relationship between the two molecular machines. To test for evolutionary conservation in human cells, we endogenously tagged human EMC5 with mNeongreen in HEK293T cells and performed an IP-MS experiment (Fig. 4a). Consistent with our findings in yeast, ATP13A1 (the human ortholog of yeast Spf1) was a main interactor of the human EMC (hEMC) (Fig. 4a, Supplementary Table 3).

**Figure 4.**
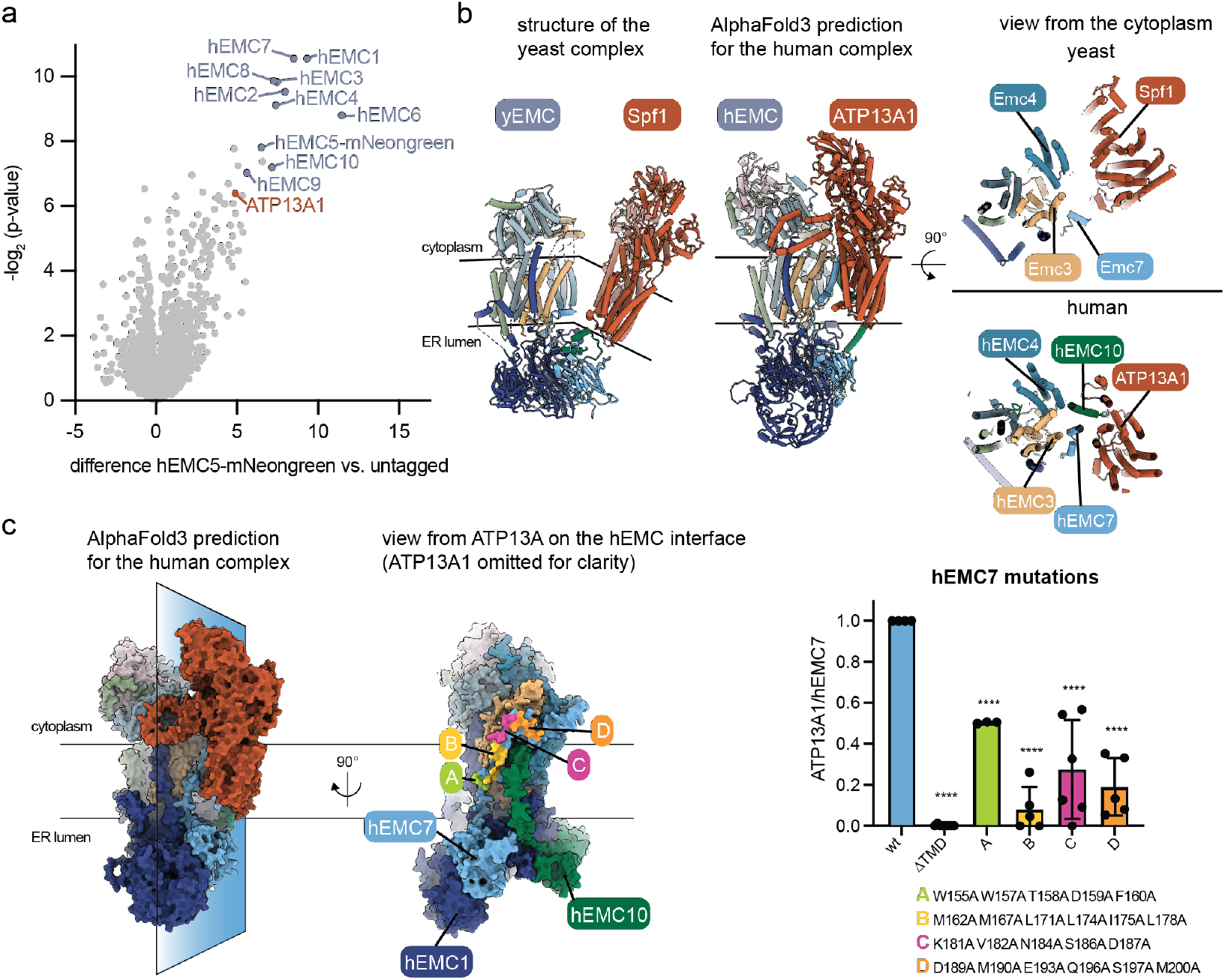
The EMC:Spf1 complex is conserved in human cells. **(A)** Interactome of endogenously tagged hEMC5-mNeongreen in HEK293T cells reveals ATP13A1, the human Spf1 ortholog, as a major interaction partner. **(B)** The experimentally obtained structure of the yeast EMC:Spf1 complex (left) in comparison to a model of the hEMC:ATP13A1 complex generated by AlphaFold 3 (middle) shows architectural conservation of the complex across species. A view of the complex from the cytoplasm (right) depicts the arrangement of the transmembrane domains in the yeast (top) and human (bottom) complex inside the lipid bilayer and shows the proximity of the transmembrane domains of hEMC7 and hEMC10 to ATP13A1. **(C)** Structural model of the hEMC:ATP13A1 complex as in b but shown in a space filling representation. A section through the model was taken and the side of the hEMC facing ATP13A1 is shown on the right. Mutations introduced into hEMC7 are highlighted in the structural model (A-D) and their effect on endogenous ATP13A1 binding was assessed by co-IP in HEK293T cells. Binding experiments were performed in at least three independent replicates (n=8 for ΔTMD, n=3 for A, n=5 for B, C, D) and normalized to the respective wt. (mean ± standard deviation, unpaired Student’s t-test * p < 0.05, ** p < 0.01, *** p < 0.005, **** p < 0.0001). Structural details and effects on hEMC assembly are shown in Extended Data Fig. 6d.

To obtain a structural model of the human EMC:ATP13A1 supercomplex we used AlphaFold 351 (SI Fig. 4). The prediction placed the hEMC and ATP13A1 in a similar orientation towards each other as observed for the yeast complex, with the insertase and the dislocase cavities facing each other (Fig. 4b, SI Fig. 4). The major interaction interface observed in the yeast complex, the lumenal dock, was partially conserved in the human model (Fig. 4b) with hEMC10 contacting ATP13A1 from the lumenal side in a similar fashion as in the yeast structure.

To validate the predicted lumenal interaction, we replaced the charged residues in this lumenal segment of hEMC10 (E172-K195) with neutral amino acids (SI Fig. 5b, 6b, c). Consistent with the model, this disrupted the interaction of hEMC with ATP13A1 while only mildly affecting overall EMC assembly (Extended Data Fig. 6b).

We next investigated the lumenal loop region of Emc7, which was critical for the yeast EMC:Spf1 interaction (Fig. 3c). This loop is shorter in hEMC7 (SI Fig. 5a, 6a) and was not predicted to contact ATP13A1 in the AlphaFold model. Indeed, comprehensive mutagenesis of the loop in hEMC7 as well as of the complementary interface in ATP13A1, corresponding to the highly effective Spf1 E1087A mutant in yeast, did not affect complex formation, suggesting that this specific interface was not required in human cells (Extended Data Fig. 6c).

These results may suggest that whereas the overall complex structure is conserved, partially different interfaces may have evolved that stabilize it: the lumenal dock seems to be less important in the human context, yet the structural prediction suggests an extended intramembrane interface between the hEMC and ATP13A1 (Fig. 4b, right). This predicted intramembrane interface involves regions of the hEMC that have not been resolved in experimental structures, namely the transmembrane domains of hEMC7 and hEMC10. Thus, to first validate the predicted positioning of these TMDs experimentally, we introduced cysteine pairs into the TMDs of hEMC7 and hEMC10 and performed in-cell crosslinking with the bismaleimide-based crosslinker BMOE. The observed crosslinks were fully consistent with the predicted direct interaction between the TMDs of hEMC7 and hEMC10 (Extended Data Fig. 7) as well as experimental evidence from other studies for the positioning of the TMD of hEMC7^1,39^. Building on this validation of the hEMC model, we next analyzed the predicted hEMC:ATP13A1 intramembrane interface in more detail. This interface mainly involves the transmembrane regions of hEMC7 (Fig. 4c, Extended Data Fig. 6d). First, we deleted the entire TMD and C-terminal region of hEMC7 (Δ159-242), resulting in a complete loss of interaction between the hEMC and ATP13A1 and, unexpectedly, in a slightly stronger association of this hEMC7 variant with the overall hEMC (Fig. 4c and Extended Data Fig. 6d). A deletion of the equivalent Emc7 TMD in yeast also resulted in reduced Spf1 binding but in contrast to the human complex negatively impacted the overall EMC assembly (Extended Data Fig. 5b). To dissect this hEMC7:ATP13A1 intramembrane interface in more detail, we introduced mutations into the individual segments of the hEMC7 TMD and C-terminus and assessed their effect on interaction with ATP13A1 as well as hEMC assembly (Fig. 4c and Extended Data Fig. 6d). This analysis further supported our structural model and defined multiple contact regions between hEMC7 and ATP13A1 within the transmembrane- and cytoplasmic regions that contributed to complex stability (Fig. 4c and Extended Data Fig. 6d). In ATP13A1, a reciprocal mutation at the border between the membrane region and the ER lumen was introduced which reduced interaction with the hEMC to half (Extended Data Fig. 6e). This mutation was also transferred to the yeast Spf1 and assessed in the recombinant insect cell setup, showing reduced complex formation (Extended Data Fig. 6f). Together, these findings indicate that the yeast EMC:Spf1 complex is conserved across spec ies, to human cells (hEMC:ATP13A1) and show roles for the same subunits (Emc7 and Emc10) in mediating the interaction.

### The Spf1 ATPase drives major conformational changes in the EMC:Spf1 complex

Spf1 is a P-type ATPase that follows the Post-Albers cycle for substrate translocation, cycling between two major states: E1 and E2. In the E1 state, the substrate binding cleft is accessible from the cytoplasm, whereas in the E2 state it faces the ER lumenal side^26,31^. To investigate the potential for Spf1 to adopt distinct functional conformations in the supercomplex with EMC, we determined its structure in the presence of the slowly hydrolysable ATP analog ATPγS to a global resolution of 3.8 Å (SI Fig. 7) and obtained a low-resolution structure in the presence of the ATP hydrolysis transition state analog AlF_4_^-^ (global resolution 6.8 Å) (SI Fig. 8). Biochemical trapping of the complex in the E2 state, or intermediates of such using other nucleotide analogs did not yield a sample that allowed structure determination by cryo-EM.

The EMC:Spf1^ATPγS^ complex was stabilized in the E1-ATP state (RMSD to 6XMQ: 0.927 Å), whereas the presence of AlF_4_^-^ and ADP most likely mimicked the E1-P state, as reported previously^26^ (Fig 5a). While the overall architecture is conserved across different states, distinct structural characteristics distinguish the different nucleotide states. In the EMC:Spf1^ATPγS^ state we observed well-defined density for nearly the entire sequence of the Spf1 “arm” – an insertion in the P-domain (L834-G956) exclusive to P5A-ATPases. Density for this arm, as well as the cytoplasmic helices of Emc3 that are contacting the arm in the EMC:Spf1^ATPγS^ complex, was completely absent in the EMC:Spf1^AlF4-^ complex. Between these two states, Spf1 moves substantially, repositioning its whole transmembrane core and cytoplasmic domains 10 Å closer towards the EMC after ATP hydrolysis (Fig. 5b). Of note, we also observed substantial conformational heterogeneity of Spf1 in our map of the endogenous EMC:Spf1 complex (Extended Data Fig. 3e).

**Figure 5.**
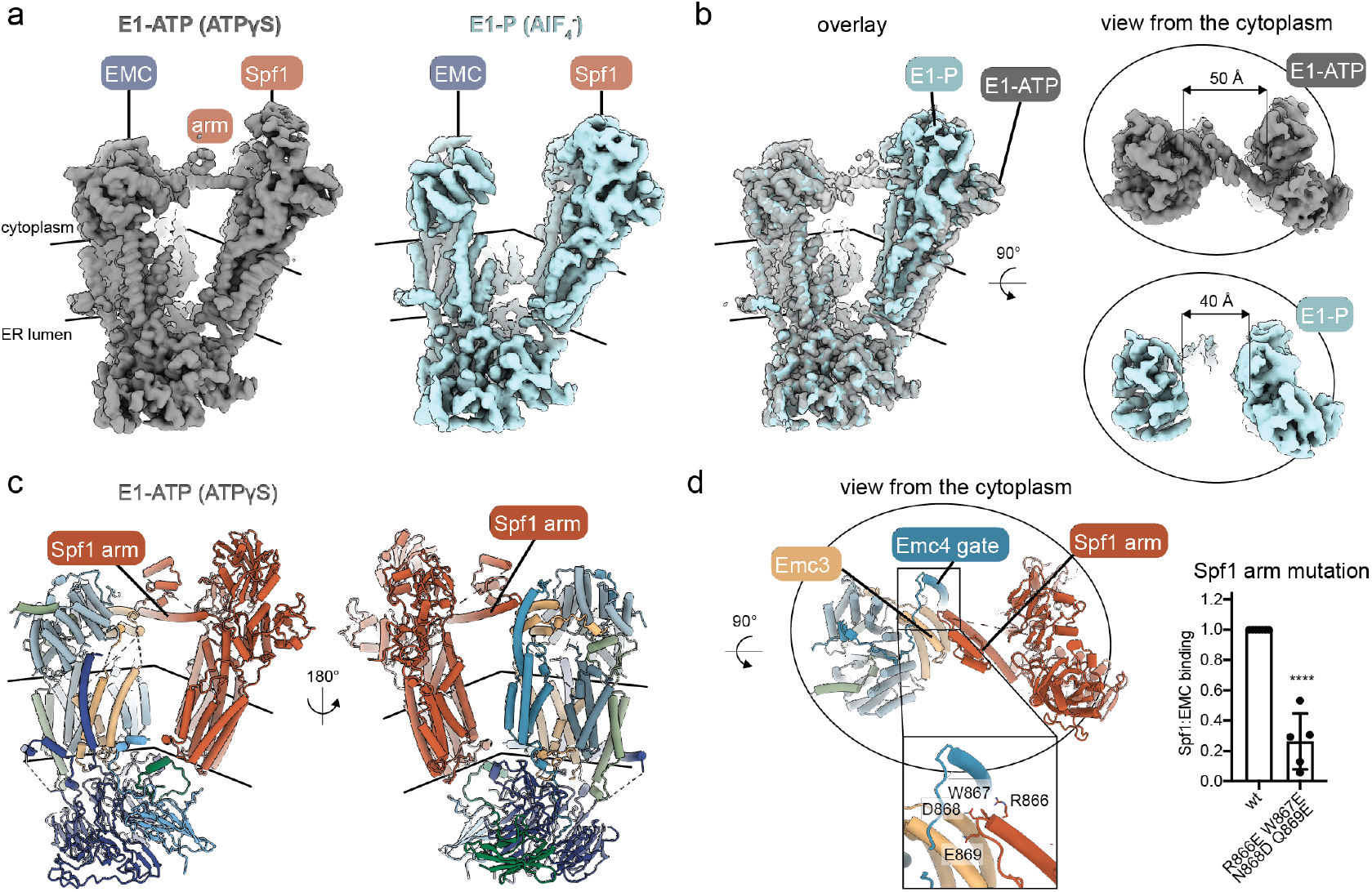
Nucleotide binding and hydrolysis by the P-type ATPase Spf1 remodels the EMC:Spf1 complex and controls access to the composite insertase-dislocase intramembrane cavity. **(A)** Cryo-EM maps of the EMC:Spf1 complex bound to the nucleotide analogs ATPγS (left, grey) AlF_4_^-^ (right, aquamarine), mimicking the E1-ATP and the E1-P state, respectively. In the E1-ATP state, the arm domain of Spf1 is visible, extending from Spf1 through the cytoplasm to the cytoplasmic cap of the EMC. Approximate boundaries of the detergent micelle are indicated with black lines. **(B)** Overlay and direct comparison of the two different nucleotide states shows how the arm domain of Spf1 can function as a lid in the E1-ATP state, blocking entry from the cytoplasm towards the composite intramembrane cavity of the complex. In the E1-P state no density for the arm domain can be observed and the size of the cavity is reduced by 10 Å due to repositioning of Spf1 towards the EMC within the micelle. Approximate boundaries of the detergent micelle are indicated with black lines. **(C)** The EMC:Spf1^ATPγS^ structure from two different views shows the arm domain of Spf1, which contacts the EMC at the cytosolic cap between Emc3 and Emc4 and blocks access to the shared substrate cavity. **(D)** Top view from the cytoplasm on the arm of Spf1 (cartoon representation), contacting the EMC. The close-up shows the tip of the arm where mutated residues are highlighted. Binding of the Spf1 tip mutation to the EMC was assessed in the recombinant insect cell system in five independent replicates and normalized to the respective wt. (mean ± standard deviation, unpaired Student’s t-test **** p < 0.0001).

The Spf1 arm itself forms an extended helical protrusion with a globular domain at its tip. In our EMC:Spf1^ATPγS^ structure it spans the cytoplasmic space between Spf1 and the EMC, contacting Emc3 at the cytoplasmic cap and the Emc4 gate (Fig. 5c, d). Previous structures of Spf1 in the E1-ATP state had resolved the arm (or plug) domain at a low resolution^26,31^ (Extended Data Fig. 8). While in these structures the arm domain points towards the detergent micelle, in our map the arm is well resolved and demonstrates another interaction site for the EMC. Notably, we observe significantly more density for Emc3 and Emc4, contacting the arm, in the presence ATPγS in comparison to the apo map.

To corroborate this nucleotide-dependent interaction interface, we placed mutations at the tip of the Spf1 arm to disrupt the interaction with Emc3 and 4. Emc3 contains a negatively charged patch at the interaction site and introduction of negative charges into the tip of the Spf1 arm domain led to significant destabilization of the complex, potentially due to electrostatic repulsion (Fig. 5d). Together, our data show that the EMC:Spf1 complex undergoes major conformational changes along the Spf1 ATPase cycle and that the enigmatic Spf1 arm appears to play a thus far unappreciated regulatory role in the EMC:Spf1 complex.

## Discussion

Our study shows that the ER of eukaryotic cells contains a high abundance supercomplex between the EMC, a multifunctional molecular machine hosting a TMD insertase^1,14,15,41^, and Spf1, an ATP-dependent transmembrane dislocase^26,52^. We determined cryo-EM structures of endogenous and recombinant EMC:Spf1 complexes which together reveal key features of this integrated machinery: i. In the EMC:Spf1 complex, the TMD insertase and dislocase cavities are juxtaposed to form a composite cavity in the membrane that could accommodate single or multiple co-client TMDs; ii. Emc7 and Emc10 together act as a recruitment platform for Spf1, providing a biochemical function for ill-characterized Emc10; iii. nucleotide-state dependent conformational cycling of Spf1 remodels the supercomplex and controls access to the composite cavity; iv. experimentally validated structural modeling shows this supercomplex and its architecture to be conserved from yeast to humans. Together, our study suggests a mechanistic framework for a joint insertase-dislocase activity in the ER membrane that surveils eukaryotic membrane protein insertion and topogenesis (Fig. 6).

**Figure 6.**
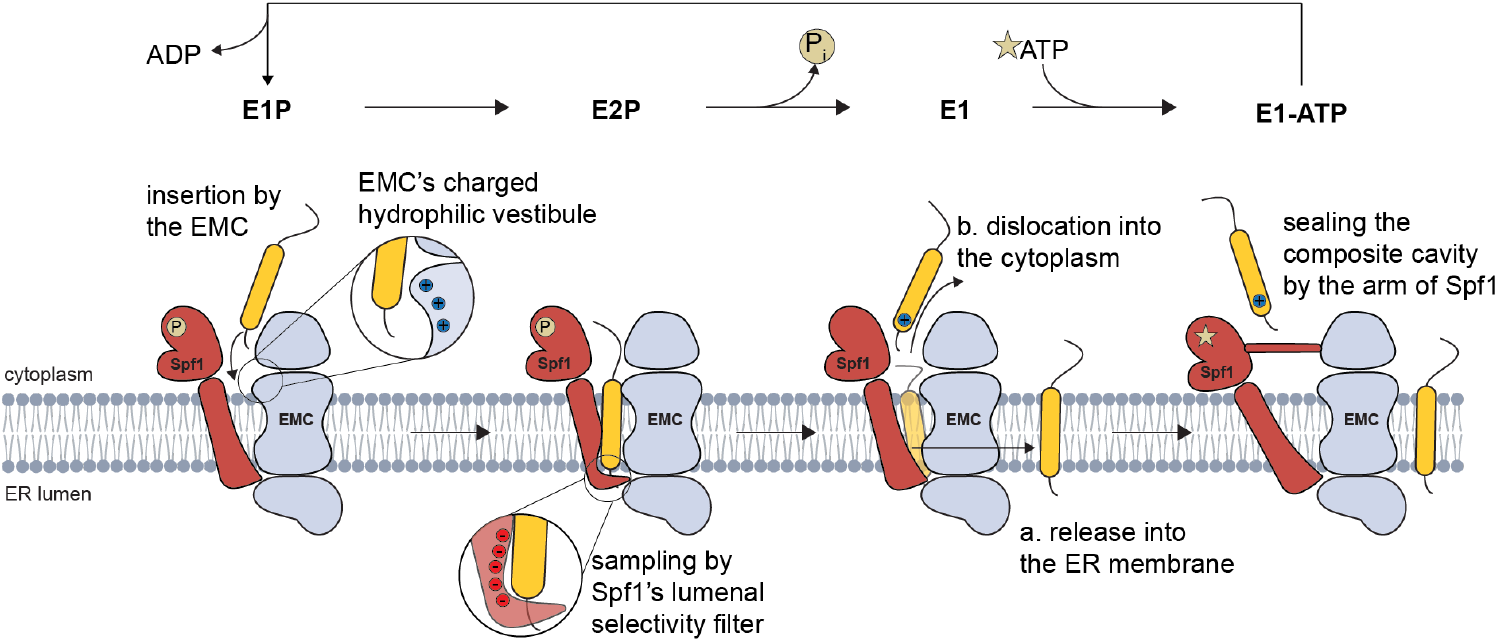
Model for an ATP-driven substrate translocation cycle by the EMC:Spf1 complex. The composite transmembrane cavity of the EMC:Spf1 complex in its E1P conformation is well accessible from the cytoplasmic side and allows the EMC to engage substrates, probe them with its selectivity filter^1^, and insert them into the lipid bilayer. In the E2P state, the substrate binding site is facing the ER lumen. Spf1’s intramembrane cleft is rich in negative charges which interact with the lumen-proximal end of the substrate TMD. If no binding occurs, the substrate may be released into the ER membrane (a.). If the lumen-proximal part of the substrate is positively charged, it may bind to Spf1 and induce a conformational shift back into the E1 conformation with the substrate binding site facing the cytosol and expelling a phosphate and the substrate (b.). Subsequent ATP binding stabilizes the Spf1 arm domain in a conformation that blocks the composite cavity and hence prevents immediate re-insertion. Note that the shared EMC:Spf1 cavity is large enough to also accommodate multiple TMDs which may suggest an action also on clients beyond individual TMDs.

### A composite cavity for membrane protein insertion and extraction

Our structural analyses reveal a shared cavity formed jointly by EMC and Spf1, which suggests that the biochemical functions of TMD insertion and extraction could be coordinated or integrated. This composite vestibule provides a structural framework for understanding TMD insertion, quality control, and potential proofreading. Both EMC and Spf1 employ selectivity filters based on charged residues. Insertion by EMC is gated at the cytosolic cavity entrance, where positive residues act as a charge-selective barrier, excluding misoriented or mismatched TMDs^1^. Conversely, Spf1 recognizes mislocalized or misoriented substrates that contain similar positively charged motifs exposed to the ER lumen^26^. This shared logic of charge-based discrimination may enable synergistic proofreading on both sides of the ER membrane during membrane protein biogenesis. Of note, the size and shape of the composite cavity suggest capacity to accommodate multipass membrane proteins. This is consistent with Spf1’s known role in extracting TMDs from complex substrates and mis-inserted signal sequences to aid correct targeting and topogenesis^25,35,53,54^. We propose that the close coupling of insertase and chaperone functions by the EMC and dislocase functions by Spf1 in this complex enhances the fidelity and efficiency of such salvage pathways.

Beyond the dimension of this cavity, the architecture of the EMC:Spf1 complex brings into close proximity parts of both proteins subject to dynamic conformational changes: the EMC insertase cavity gate, comprising the TMDs of Emc4, and the Spf1/ATP13A1 client helix binding pocket, which is remodeled by the ATPase cycle. Excitingly, we observe the Emc4 gate helices to come as close as ∼5Å to Spf1 at the lumenal border of the transmembrane region and to be contacted by the arm domain of Spf1 in the cytoplasm, suggesting that previously observed conformational changes observed in the Emc4 gate^38^ may be coupled to Spf1’s ATP-dependent conformational cycling.

### Spf1 constitutes an ATPase-driven molecular machine in the supercomplex

An important mechanistic implication from our structural investigations of the EMC:Spf1 complex is the conformational coupling between the ATPase cycle of Spf1 and the EMC’s insertase activity. This provides insights into a key question in the field: are membrane protein insertases/chaperones energy-dependent and if so, how is ATP hydrolysis functionally coupled to insertion? Our structures show pronounced conformational changes in the EMC:Spf1 complex that are driven by the ATPase cycle of Spf1 and regulate access to the EMC insertase cavity. Specifically, the unique Spf1 arm domain which interacts directly with two cytoplasmic helices of Emc3 and the extension of the first helix of the Emc4 gate in the ATP-bound state. In this conformation the arm occludes the entrance to the composite cavity from the cytoplasm. A previous study on *Chaetomium thermophilum* Spf1 speculated about a similar function for the arm domain and could resolve dislocation from the cavity entrance upon ATP-hydrolysis, hence functioning as a dynamic “plug”^31^. Based on these collective findings, we propose that this ATP-regulated shielding may function as gating mechanism for the overall EMC:Spf1 complex and may be important for avoiding re-integration of freshly extracted TMDs.

In general, ATP hydrolysis is a hallmark of high-fidelity molecular machines, often employed to provide directionality or proofread critical steps. Our data now shows that such an energy-coupled regulation could also be used for controlling protein insertion by the EMC in the ER.

### The EMC as a modular membrane protein quality control hub

In recent years, three distinct complexes of the human EMC have been structurally characterized (Extended Data Fig. 9), revealing key aspects of its functions in membrane protein biogenesis: the structure of a bound assembly intermediate of a voltage gated anion channel^18^ provided strong evidence for a holdase function of the EMC^19^; the EMC:BOS holo-insertase complex introduced the concept of the EMC as a central hub in the ER which orchestrates membrane protein biogenesis^13^; and a structure of the EMC bound to endogenous VDAC^55^ raised new questions on further roles of the EMC in inter-organelle communication. The EMC:Spf1 structure now adds another distinct mode of interaction in comparison to these human EMC complexes, mediated by a so far largely uncharacterized interaction interface involving the subunits Emc7 and Emc10. This significantly extends the view of the EMC as a central hub for membrane protein quality control.

Taken together, our results identify the EMC:Spf1 complex as an evolutionarily conserved, energy-dependent insertase– dislocase machine in the ER of eukaryotic cells. It integrates directional insertion, extraction, and error-correction within a single structural entity, establishing a new paradigm for membrane protein quality control in the ER. Furthermore, our study reveals that the EMC engages clients and interaction partners via distinct interfaces. It will now be key to analyze how engagement of the different sites is regulated to ultimately govern the fate of membrane protein clients.

## Methods

### DNA constructs and molecular biology

DNA for all yeast genes (yEmc1, yEmc2, yEmc3, yEmc4, yEmc5, yEmc6, SOP4, yEmc10, Spf1) was isolated from genomic DNA of *S. cerevisiae* BY4741 and epitope tags (FLAG, TwinStrepII) were introduced via PCR, before cloning into a pLIB vector and ultimately combining all EMC subunits into one expression vector using the biGBac multi-gene cloning method^56^. DNA for human ATP13A1, hEMC7 and hEMC10 was obtained from the NIH Mammalian Gene Collection cDNA library and cloned into respective vectors for expression in mammalian cells (pEG, pCDNA3.4). The pSpCas9(BB)-2A-Puro (PX459) vector was obtained from Addgene (#48139). The gene for mNeongreen used for endogenous tagging was a kind gift of Marius Lemberg, University of Cologne. All constructs were derived from these using standard molecular biology procedures and Gibson assembly and verified by DNA sequencing.

### Yeast strains

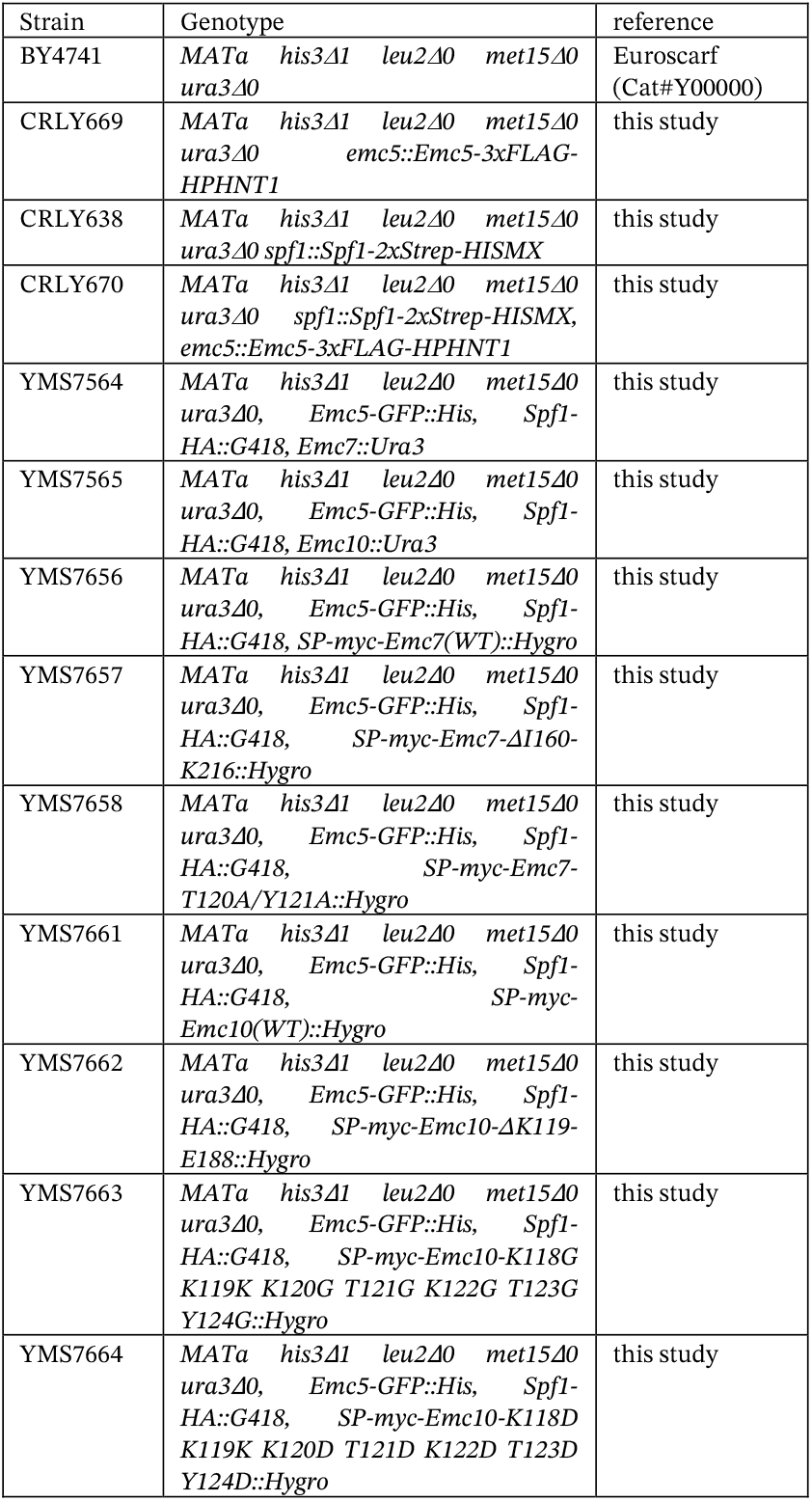

| Strain | Genotype | reference |
| --- | --- | --- |
| BY4741 | *MATa his3Δ1 leu2Δ0 met15Δ0 ura3Δ0* | Euroscarf (Cat#Y00000) |
| CRLY669 | *MATa his3Δ1 leu2Δ0 met15Δ0 ura3Δ0 emc5::Emc5-3xFLAG-HPHNT1* | this study |
| CRLY638 | *MATa his3Δ1 leu2Δ0 met15Δ0 ura3Δ0 spf1::Spf1-2xStrep-HISMx* | this study |
| CRLY670 | *MATa his3Δ1 leu2Δ0 met15Δ0 ura3Δ0 spf1::Spf1-2xStrep-HISMx, emc5::Emc5-3xFLAG-HPHNT1* | this study |
| YMS7564 | *MATa his3Δ1 leu2Δ0 met15Δ0 ura3Δ0, Emc5-GFP::His, Spf1-HA::G418, Emc7::Ura3* | this study |
| YMS7565 | *MATa his3Δ1 leu2Δ0 met15Δ0 ura3Δ0, Emc5-GFP::His, Spf1-HA::G418, Emc10::Ura3* | this study |
| YMS7656 | *MATa his3Δ1 leu2Δ0 met15Δ0 ura3Δ0, Emc5-GFP::His, Spf1-HA::G418, SP-myc-Emc7(WT)::Hygro* | this study |
| YMS7657 | *MATa his3Δ1 leu2Δ0 met15Δ0 ura3Δ0, Emc5-GFP::His, Spf1-HA::G418, SP-myc-Emc7-ΔI160-K216::Hygro* | this study |
| YMS7658 | *MATa his3Δ1 leu2Δ0 met15Δ0 ura3Δ0, Emc5-GFP::His, Spf1-HA::G418, SP-myc-Emc7-T120A/Y121A::Hygro* | this study |
| YMS7661 | *MATa his3Δ1 leu2Δ0 met15Δ0 ura3Δ0, Emc5-GFP::His, Spf1-HA::G418, SP-myc-Emc10(WT)::Hygro* | this study |
| YMS7662 | *MATa his3Δ1 leu2Δ0 met15Δ0 ura3Δ0, Emc5-GFP::His, Spf1-HA::G418, SP-myc-Emc10-ΔK119-E188::Hygro* | this study |
| YMS7663 | *MATa his3Δ1 leu2Δ0 met15Δ0 ura3Δ0, Emc5-GFP::His, Spf1-HA::G418, SP-myc-Emc10-K118G K119K K120G T121G K122G T123G Y124G::Hygro* | this study |
| YMS7664 | *MATa his3Δ1 leu2Δ0 met15Δ0 ura3Δ0, Emc5-GFP::His, Spf1-HA::G418, SP-myc-Emc10-K118D K119K K120D T121D K122D T123D Y124D::Hygro* | this study |

C-terminal tagging of Emc5 and Spf1 in *S. cerevisiae* for initial IP-MS and structural studies was achieved using standard techniques^57,58^ and strains were verified by sequencing and western blotting.

Yeast strains for co-immunoprecipitation studies were constructed using the lithium acetate-based transformation protocol^59^. For introducing *emc7* and *emc10* variants into their endogenous loci, YMS7564 and YMS7565 were used respectively. After transformation with PCR products encoding the required variants and a Hygromycin (Hygro) cassette, cells were plated on solid agar media containing 5-FOA (5-fluoro-orotic acid monohydrate) for counter-selection, and Hygro for positive selection. Strains were validated by PCR, followed by genomic sequencing of either the *emc7* or *emc10* locus, to ensure that the inserted variants were in-frame with their endogenous 5’ UTR regions.

### Human cell lines

HEK293T cells (Sigma Aldrich 12022001) were cultured in Dulbecco’s Modified Eagle Medium (DMEM) high glucose supplemented with 10% (v/v) fetal bovine serum (FBS; Gibco) and 1% (v/v) antibiotic-antimycotic supplement (Gibco; final concentration 100 units/mL penicillin, 0.1 mg/mL streptomycin and 2.5 µg/mL Amphotericin B) at 37 °C with 5% CO_2_.

### Detergents

Digitonin (Sigma Aldrich) was further purified as described previously^60^. In brief, 30 g digitonin was dissolved in 500 mL boiling double-distilled water, kept at 4 °C for 36 hours before the precipitate was removed by centrifugation at 20,000 xg for 15 min. 10 mM Tris-HCl was added to adjust the pH to 8.0. The solution was finally passed sequentially through Q-Sepharose and S-Sepharose resins by gravity flow and stored as 5 % stock solution at -20 °C. GDN (Anatrace) was dissolved in double-distilled water at 5% (w/v) and stored at -20 °C.

### Isolation of endogenous EMC and Spf1 from *S. cerevisiae*

Microsome preparation and immunoprecipitations were performed as described previously^17^. *S. cerevisae* were grown in YPD until late log phase, harvested and resuspended in 1 mL lysis buffer (50 mM HEPES-KOH pH 7.5, 150 mM KoAc, 2 mM MgOAc, 1 mM CaCl_2_, 0.2 M Sorbitol, 2X complete EDTA-free protease inhibitor (Roche)) per 6 grams pellet and frozen dropwise in liquid nitrogen. Cell walls were disrupted by cryo-milling using a Spex SamplePrep Freezer/Mill. The powder was thawed and, after addition of 1 mL lysis buffer per gram pellet, nutated at 4 °C for 45 min and dounced in a dounce homogenizer to dissolve in lysis buffer. The lysate was spun down twice at 1,000 xg for 10 mins to remove unbroken cells and large fragments before the supernatant was transferred to an ultracentrifuge tube and centrifuged at 50,000 xg for 20 min to pellet membranes. The microsomes were resuspended in a small volume of lysis buffer and flash frozen.

15 mL of IP buffer (50 mM HEPES-KOH pH 7.5, 150 mM KOAc, 2 mM MgOAc, 1 mM CaCl_2_, 15% (v/v) glycerol, 1x complete EDTA-free protease inhibitor) containing 2% (w/v) digitonin were added and the microsomes were nutated for at least 1 h at 4 °C to solubilize. Insoluble material was removed via centrifugation at 50,000 xg and the supernatant was added to FLAG M2 agarose beads (EMC) or Strep-Tactin agarose (IBA) for affinity isolation for 90 min. The resin was washed extensively before elution with 0.15 mg/ml 3XFLAG peptide or 5 mM Desthiobiotin. For interactomics, bound protein was not eluted but directly digested on the beads (see section mass spectrometry).

### Purification of endogenous EMC for cryo-EM

The large-scale purification of endogenously tagged EMC was performed analogous to the procedure described above. To obtain a high biomass, a yeast strain containing Emc5-3xFLAG and Spf1-TSII was grown in a fermenter to high cell density before harvesting, dropwise freezing in and cryo-milling. 700 g of milled powder were used as input and a FLAG-pulldown was performed as described above. Elution fractions were combined and concentrated before injection onto a Superose 6 column and size exclusion chromatography in SEC buffer (20 mM HEPES pH 7.5, 150 mM KOAc, 0.03% (w/v) digitonin). Fractions were analyzed by SDS-PAGE, pooled and concentrated before plunging.

### Expression and purification of recombinant EMC:Spf1 from insect cells

Bacmids were produced in DH10EMBacY *E. coli* for subsequent transfection into Sf9 insect cells (Thermo Fisher B85502) to generate baculovirus. Hi5 insect cells (BTI-TN-5B1-4; Thermo Fisher 11496016) were co-infected with respective baculoviruses and protein expression was carried out for 3 days. Cells were harvested, washed in ice-cold PBS and lysed in hypotonic lysis buffer (10 mM HEPES pH 7.5, 1 mM MgCl_2_) supplemented with complete EDTA-free protease inhibitor (Roche) and 0.001 mg/ml Benzonase. The membrane fraction was pelleted by centrifugation at 50,000 xg for 30 mins and resuspended by dounce homogenization in IP buffer (50 mM HEPES-KOH pH 7.5, 150 mM KOAc, 2 mM MgOAc, 1 mM CaCl_2_, 15% (v/v) glycerol). Membranes were solubilized by addition of purified digitonin to a final concentration of 1.5% (w/v) and rotated for 3 hours. Insoluble material was removed by centrifugation at 50,000 xg for 30 mins and the supernatant was incubated under agitation with Strep-Tactin agarose resin (IBA lifesciences) to capture the Emc5-TSII tagged complex. After washing with a total of 20 column volumes of IP buffer with 0.1% (w/v) digitonin, bound protein was eluted with IP buffer containing 5 mM D-desthiobiotin and 0.1% (w/v) digitonin. Elution fractions were analyzed via SDS-PAGE for protein content, then pooled and incubated with anti-FLAG M2 resin (Sigma) over night. Beads were washed with 20 column volumes of IP buffer with 0.1% (w/v) digitonin and bound EMC:Spf1 complexes were eluted with IP buffer supplemented with 0.15 mg/ml 3XFLAG peptide and 0.1% (w/v) digitonin. Protein containing elution fractions were pooled and concentrated to 10 mg/ml and subjected to size exclusion chromatography on a Superose 6 column in SEC buffer (20 mM HEPES pH 7.5, 150 mM KOAc, 0.03% (w/v) digitonin). All fractions were analyzed via SDS-PAGE and respective fractions were pooled and concentrated via centrifugation.

### Mass spectrometry

#### Experimental design

All samples were prepared in triplicates. Immunoprecipitations from yeast were performed as described above for isolation of the endogenous EMC for cryo-EM. Immunoprecipitations from mammalian cells were performed as described. To exclude clonal effects, triplicates of three different endogenously tagged HEK293T clones were compared to untagged HEK293T cells.

#### Sample preparation

The beads were washed 1x with PBS, the buffer was removed and the beads were incubated for 20 mins at 37°C 100 µL with SDC buffer (1% Sodium deoxycholate, SDC, Sigma-Aldrich; 40 mM 2-Chloroacetamide, CAA, Sigma-Aldrich; 10 mM Tris(2-carboxyethyl)phosphine, TCEP, Thermo Fisher Scientific in 100 mM Tris, pH 8.0). Then, the samples were diluted with 100 µL of milliQ water and the proteins were digested overnight at 37 °C by addition of 0.5 µg trypsin (Promega). The supernatant was collected with the help of a magnetic rack and was acidified with trifluoroacetic acid (TFA; Merck) to a final concentration of 1%. Precipitated SDC was removed by centrifugation and the peptide mixture was desalted via SCX StageTips. After elution, the samples were vacuum dried and dissolved in 15 µL of buffer A (0.1% formic acid). The peptide mixture was acidified with TFA to a final concentration of 1%, followed by desalting of the peptides via SCX StageTips. Samples were vacuum dried and re-suspended in 6 µl of buffer A (0.1% formic acid).

#### LC-MS/MS data acquisition

LC-MS/MS data acquisition was performed using an Easy nanoLC coupled to a Q Exactive HF mass spectrometer (Thermo Fisher) or a timsTOF Pro mass spectrometer (Bruker Daltonic). Peptides (3 µl, corresponding to 500 or 200 ng) were loaded onto a 30-cm column (inner diameter: 75 microns; packed in-house with ReproSil-Pur C18-AQ 1.9 micron beads, Dr. Maisch GmbH) via the autosampler of the Thermo Easy-nLC 1200 (Thermo Fisher Scientific). The column was heated to 60°C. Peptides were separated with a flow rate of 250 nL/min by a gradient of buffer B (80% ACN, 0.1% formic acid) from 2% to 30%B and to 60%B over 10 minutes then to 95%B over the next 5 minutes and finally the percentage of buffer B was maintained at 95%B for another 5 minutes.

The QExactive HF mass spectrometer was operated in a data-dependent mode with MS1 scans from 300 to 1750 m/z (resolution of 60000 at m/z =200), and up to 15 of the top precursors were selected for fragmentation using higher energy collisional dissociation (HCD with a normalized collision energy of value of 28). The MS2 spectra were recorded at a resolution of 15000 (at m/z = 200). AGC target for MS1 and MS2 scans were set to 3×106 and 1×105 respectively within a maximum injection time of 100 and 25 ms for MS1 and MS2 scans respectively.

Data acquisition on the timsTOF Pro was performed using otofControl 6.0. The mass spectrometer was operated in data-dependent PASEF mode with one survey TIMS-MS and ten PASEF MS/MS scans per acquisition cycle. Analysis was performed in a mass scan range from 100-1700 m/z and an ion mobility range from 1/K0 = 1.6 Vs cm^-2^ to 0.6 Vs cm^-2^ using equal ion accumulation and ramp time in the dual TIMS analyzer of 100 ms each at a spectra rate of 9.52 Hz. Suitable precursor ions for MS/MS analysis were isolated in a window of 2 Th for m/z < 700 and 3 Th for m/z > 700 by rapidly switching the quadrupole position in sync with the elution of precursors from the TIMS device. The collision energy was lowered stepwise as a function of increasing ion mobility, starting from 52 eV for 0-19% of the TIMS ramp time, 47 eV from 19-38%, 42 eV from 38-57% 37 eV from 57-76% and 32 eV until the end. We made use of the m/z and ion mobility information to exclude singly charged precursor ions with a polygon filter mask and further used ‘dynamic exclusion’ to avoid re-sequencing of precursors that reached a ‘target value’ of 20,000 a.u. The ion mobility dimension was calibrated linearly using three ions from the Agilent ESI LC/MS tuning mix (m/z, 1/K0: 622.0289, 0.9848 Vs cm^-2^; 922.0097, 1.1895 Vs cm^-2^; 1221.9906, 1.3820 Vs cm^2^).

### Data Analysis

Raw data were processed using the MaxQuant computational platform (version 2.0.1.0)^61^ with standard settings applied. Shortly, the peak list was searched against the UniProt database of *S. cerevisiae* and the bait protein. Cysteine carbamidomethylation was set as static modification, and methionine oxidation and N-terminal acetylation as variable modifications. The match-between-run option was enabled, and proteins were quantified across samples using the label-free quantification algorithm in MaxQuant generating label-free quantification (LFQ) intensities.

MaxQuant outputs were imported into Perseus version 1.6.15.0^62^. Potential contaminants, reverse identified hits and proteins only identified by site were removed by filtering before LFQ intensities were log_2_ transformed, rows with less than three valid values in at least one group were removed before missing values were imputed from a downshifted normal distribution. Samples were compared by two sample student’s t-tests. Classifications shown in SI Figure 1 were derived from the available Uniprot annotations and gene ontology terms.

### Single particle cryo-EM

#### Sample preparation

The EMC:Spf1 apo sample was concentrated to 5.0 mg/ml, the endogenous yeast EMC sample was concentrated to 1.4 mg/ml. For the EMC:Spf1^ATPγS^ sample, the purified protein was concentrated to 4.5 mg/ml and pre-incubated with 2 mM ATPγS (Jena Bioscience) for 30 min on ice. For the EMC:Spf1^AlF4-^ sample, the purified protein was concentrated to 4.5 mg/ml, pre-incubated with 1.7 mM ADP and 10 mM NaF for 10 min on ice, before adding 2 mM of AlF_3_ and incubation for 1 h on ice. All samples were passed through a centrifugal filter to remove potential aggregates. 3.6 µl of freshly purified, concentrated sample was applied on glow-discharged R1.2/1.3 300 mesh holey carbon grids (Quantifoil) and plunge-frozen in liquid ethane on a Vitrobot Mark IV (Thermo Fisher; blot force 4, blot time 3 s, 4 °C, 100% humidity).

#### Data collection

The endogenous yeast EMC grids were screened and data was collected using SerialEM v3.8^63^ on a Glacios cryo-TEM (FEI) operated at 200 kV and equipped with a K2 Summit direct electron detector (Gatan). A total of 3,690 micrographs were collected at a 22,000x magnification, corresponding to a pixel size of 1.885 Å/px, in a defocus range of -1.0 to - 2.6 µm, with a total dose of 60 e^-^/Å^2^ fractionated over 40 frames.

All grids of the recombinant EMC:Spf1 sample were screened using SerialEM v4.1^63^ on a Talos Arctica cryo-TEM (FEI) operated at 200 kV and equipped with a K3 direct electron detector (Gatan). Screening datasets and the final dataset for EMC:Spf1^AlF4-^ were collected at a 22,000x magnification, corresponding to a pixelsize of 1.841 Å/px, in a defocus range of -1.0 to -2.6 µm, with a total dose of 60 e^-^/Å^2^ fractionated over 40 frames.

High-resolution datasets of the recombinant EMC:Spf1 apo and incubated with ATPγS were collected using SerialEM v4.1^63^ on a Titan Krios cryo-TEM (FEI) operated at 300 kV, equipped with a K3 direct electron detector (Gatan) in counting mode and a Bio Quantum-LS post-column energy filter (Gatan; 10 eV). The datasets were recorded at a 105,000x magnification, corresponding to a pixel size of 0.8512 Å/px, in a defocus range of -0.7 and -2.2 µm, with a total dose of 65.1 e^-^/Å^2^ (apo) and 62.62 e^-^/Å^2^ (ATPγS).

### Data processing

#### Recombinant EMC:Spf1 complex (SI Fig. 2)

Following import into RELION 5.0^64,65^, the micrographs underwent motion correction and dose-weighting, leveraging RELION’s built-in MotionCorr2 implementation. Contrast transfer function (CTF) estimation was then performed with CTFFIND version 4.1.9^66^.

For the apo dataset, we initially picked 3,817,633 particles using Gautomatch^67^. These were then subjected to 3D classification on a binned dataset (3.05 Å pixel size) with a regularization parameter of T=16, yielding 924,465 particles for further 3D classification. After multiple rounds of 3D classification and orientation refinements, the particle count was narrowed to 645,091. These particles were then extracted to full pixel size for additional 3D refinements. We employed RELION’s multibody refinement implementation to resolve the EMC and Spf1 components. Separate post-processing of these two bodies resulted in resolutions of 3.8 Å for EMC and 4.2 Å for Spf1. We used DeepEMhancer^68^ to sharpen the maps, and a composite map was constructed using PHENIX^69^ with the DeepEMhancer sharpened EMC and Spf1 maps as input.

#### Endogenous yeast EMC (SI Fig. 3)

Raw movies were processed in CryoSPARC v.4.7.1^70^ using motion correction and patch CTF estimation and manually curated, excluding 429 exposures. Templates for particle picking were generated based on manual picking and initial 3D model generation and resulted in 814,288 particles that were extracted at a binned size (6.7 Å pixel size). These were subjected to one round of 2D classification and one round of heterogeneous refinement based on the initial 3D models. The handedness of the best class was flipped before subjecting it to one round of non-uniform refinement. The particles were extracted at full pixel size (1.885 Å) and underwent further refinement before being used to train a 3D Flex model^71^ with 4 latent modes. 80,000 particles were used to reconstruct the flexible volume which was further refined after reference-based motion correction, resulting in a consensus map of 8.7 Å resolution. A mask around the EMC allowed to locally refine the EMC to 8.6 Å resolution, resolving all its transmembrane helices.

#### EMC:Spf1^ATPγS^ (SI Fig. 7)

Following import into RELION 5.0, the micrographs underwent motion correction and dose-weighting, using RELION’s built-in MotionCorr2 implementation. CTF estimation was then performed with CTFFIND version 4.1.9.

For the ATPγS dataset, 2,275,267 particles were initially picked using Gautomatch. These were subjected to 3D classification on a binned dataset (3.05 Å pixel size) with a regularization parameter of T=16, yielding 514,084 particles. Multiple rounds of classification and refinement were carried out to discard poor-quality particles, resulting in a final subset of 33,661 high-quality particles. These particles were re-extracted at full pixel size for subsequent refinement and postprocessing, ultimately yielding a 3.8 Å reconstruction.

#### EMC:Spf1^AlF4-^ (SI Fig. 8)

Raw movies were processed in CryoSPARC v.4.7.1^70^ using motion correction and patch CTF estimation. Particles were first picked using the blob picker tool to create an initial model for 2D template generation and template-based picking of 2,059,914 particles. After two rounds of 2D classification, 228,528 particles were used to generate initial 3D models which served as input models for two rounds of heterogeneous refinement of the full particle set. The best class was refined once, extracted at full pixel size (1.885 Å) and the subjected to two further rounds of refinement. After an additional round of 3D classification, the best class consisting of 103,225 particles was refined and post-processed, resulting in a final global resolution of 6.8 Å. DeepEMhancer^68^ was used to further sharpen the final map.

#### Model building

The 3.8 Å reconstruction map of the ATPγS-bound complex revealed well-defined main-chain and side-chain densities, enabling the fitting of a previously determined structure for EMC (PDB: 7KRA) and a model for Spf1 predicted by AlphaFold^72,73^ (AF-P39986-F1). Initial docking was performed using UCSF Chimera^74^, allowing each domain to move independently during rigid-body refinement in PHENIX^75^. Model building and real-space refinement were carried out iteratively to optimize the fit. Manual adjustments were made in COOT^76^, and real space refinements were conducted using phenix.real_space_refine.

Models for the 3.8 Å EMC and the 4.2 Å SPF1 apo structure were built using the same approach. Due to limited resolution in the nucleotide-binding domains, side-chain positioning was often ambiguous; in such cases, rotamers from previously published structures were retained when appropriate. DeepEMhancer-sharpened as well consensus refinement maps were used to guide manual model building in COOT. Real space refinements in PHENIX were done against postprocessed maps from RELION.

#### Figure preparation

All structural figures were prepared using ChimeraX-1.9^77^.

### AlphaFold structure prediction

The structural model for the human EMC-ATP13A1 complex was predicted using AlphaFold3^51^ on the AlphaFold Server. Protein sequences for the human EMC subunits 1, 2, 3, 4, 5, 6, 7, 8, 10 and ATP13A1 were derived from Uniprot using the most common isoform, if applicable the signal peptide was removed, and the following posttranslational modifications were added, as annotated in the Uniprot database:

hEMC1: N348 N-acetyl-beta-D-glucosamine

N796 N-acetyl-beta-D-glucosamine

N891 N-acetyl-beta-D-glucosamine

hEMC2: K254 N6-acetyl-L-lysine

hEMC4: T1 Phosphothreonine

hEMC10: N157 N-acetyl-beta-D-glucosamine

ATP13A1: N286 N-acetyl-beta-D-glucosamine

N419 N-acetyl-beta-D-glucosamine

S898 Phosphoserine

S904 Phosphoserine

### Co-immunoprecipitations from *S. cerevisiae*

All yeast strains were grown on agar plates containing the respective selection markers. Single colonies were used to inoculate overnight cultures in selection medium. Cultures were diluted to an OD_600_ of 0.2 in YPD medium and grown until mid-log phase was reached. 80 OD_600_ units were collected by centrifugation and cells were washed with H_2_O before flash freezing. The pellet was resuspended in 1x IP buffer with 1% (w/v) purified digitonin and complete EDTA-free protease inhibitor (Roche) and cell lysis was carried out in a FastPrep-24 benchtop homogenizer system (MP Biomedicals) using the lysing matrix C (1 mm silica spheres). The supernatant was collected after centrifugation at 16,000 xg for 10 min and incubated with anti-GFP magnetic agarose (Chromotek) for 2 h under agitation. Beads were washed with IP buffer with 0.1 % (w/v) digitonin and bound protein was eluted by addition of 2x Lämmli buffer with 10% (v/v) β-mercaptoethanol and incubation at 60 °C for 10 min.

### Co-immunoprecipitations from insect cells

Spf1 mutants and wt yeast EMC were co-expressed in Hi5 insect cells by bacoluvirus co-infection as described above. Cells were harvested 3 days post infection, pellets were washed with ice-cold PBS and lysed in 1x IP buffer supplemented with 1% (w/v) digitonin and 20 µg/ml aprotinin, 10 µg/ml leupeptin, 10 µg/ml pepstatin and 1 µg/ml Benzonase. Membranes were solubilized for 90 min before removing insoluble material by centrifugation at 20,000 xg for 30 mins. The soluble material was incubated with magnetic StrepTactin resin (PureCube HiCap StrepTactin MagBeads, Cube Biotech) for 90 mins under constant agitation. The beads were washed 5 times with 1x IP buffer supplemented with 0.01% (w/v) digitonin. Co-immunoprecipitated proteins were eluted by boiling the beads in 2x Lämmli buffer with 10% (v/v) βmercaptoethanol at 60 °C for 10 min.

### Endogenous tagging of MMGT1 (human EMC5) with mNeongreen by CRISPR/Cas9-mediated genome editing

#### Guide design and construction of the homology directed repair (HDR) template

Two single guide (sg)RNAs were designed against the C-terminus of the MMGT1 gene, targeting the sense and anti-sense DNA strand of exon 4 before and after the endogenous stop codon.

guide 1: CACCGTTTTTACAAATTATAATAAT

guide 2: CACCGTAACTTCAATGATGTGTTAG

The guides were cloned individually into the pSpCas9(BB) vector as described previously^78^ and a dual expression cassette vector was constructed by linearization via restriction digest with XbaI and insertion of the second sgRNA expression cassette via Gibson assembly. The HDR template was constructed by amplifying a 500 bp 3’ overhang and a 600 bp 5’ overhang from the endogenous MMGT1 locus adjacent to the guide cutting sites. The protospacer adjacent motifs (PAMs) were mutated to prevent recognition by the sgRNA. A 15 amino acid linker and the gene for mNeongreen were inserted between the two flanking regions. The construct was amplified via PCR, analyzed on an agarose gel and purified via a silica-membrane-based spin column (QIAquick PCR purification kit, Qiagen).

#### Transfection, antibiotic selection, FACS sorting of single cell clones

HEK293T cells were seeded 24 h prior to transfection with the dual guide and Cas9 containing plasmid and a 4-fold molar excess of the purified HDR template using Lipofectamine2000 (Thermo Fisher) as transfection reagent according to the manufacturer’s instructions. Selection for transfected cells with 1 µg/ml Puromycin (Gibco) was carried out 48 hours after transfection for 96 h, replacing the antibiotic-containing medium every 24 h. Selected cells were allowed to recover and proliferate for 5 days before the top 0.9 % mNeongreen-positive single cell clones were isolated via fluorescence activated cell sorting (FACS) on a CytoFLEX SRT instrument (Beckman Coulter).

#### Clonal validation

Individual clones were expanded and analyzed for the presence of the genomic insertion on both alleles via PCR-based screening and for the expression of the EMC5-mNeongreen fusion protein by western blot. Positive clones were further validated by deep sequencing of the insert and flanking genomic regions.

### Transient transfections

For transient transfections, 600,000 HEK293T cells were seeded in a p35 well 24 hours prior to transfection. 1 µg of DNA per construct was transfected using Metafectene Pro (Biontex) according to the manufacturer’s instructions. Protein expression was analyzed 24 hours after transfection. In-cell cysteine crosslinking Cells for this experiment were cultivated in poly-D-lysine coated dishes. Before crosslinking, cells were washed briefly with PBS before adding 250 µM BMOE (Thermo Fisher, dissolved in anhydrous DMSO to 20 mM) in PBS with 5 mM EDTA. Crosslinking was performed at room temperature in the dark for 1 h before quenching of the reaction with 20 mM DTT in PBS for 10 mins. All subsequent lysis and IP steps were performed on ice as described below, except that 1x RIPA buffer (50 mM Tris-HCl pH 7.5, 150 mM NaCl, 1% (v/v) NP-40, 0.5% (w/v) deoxycholate, 0.1.% (w/v) SDS) was used for lysis and washing.

### Co-immunoprecipitations from HEK293T cells

Cells were briefly washed twice with ice cold PBS before adding lysis buffer (50 mM Tris-HCl pH 7.5, 150 mM NaCl, 1x complete protease inhibitor EDTA-free (Roche), 0.5% (w/v) GDN). Membranes were solubilized for 20 mins on ice before centrifugation at 15,000 xg for 15 min to remove insoluble material. A small fraction of the lysate was kept as input sample and supplemented with 5x Lämmli buffer with 10% (v/v) β-mercaptoethanol to a final concentration of 1x and denatured at 60 °C for 10 mins. The remaining lysate was added to pre-coupled magnetic beads (anti-FLAG M2 magnetic beads, Millipore; Pierce anti-HA magnetic beads, Thermo Fisher; MagStrep Strep-Tactin XT beads, IBA) and incubated for 90 min. Unbound material was removed in three subsequent wash steps using wash buffer (50 mM Tris HCl pH 7.5, 150 mM NaCl, 0.01 % (w/v) GDN). For western blot analysis, immunoprecipitated protein was eluted by denaturation using 2x Lämmli buffer supplemented with 10% (v/v) β-mercaptoethanol and heating samples to 60 °C for 10 mins.

### SDS-PAGE, western blots and quantifications

Protein samples were separated on self-made 4-22% or 12 % Tris-glycine SDS-PAGE gels. For total protein analysis, gels were stained with Coomassie brilliant blue stain (homemade) or Oriole Fluorescent Gel Stain (Biorad). SDS-PAGE gel band intensities were quantified using the Fiji distribution of ImageJ^79^. In brief, the intensity profile of the gel line was plotted after performing background correction. Peaks were identified using the first derivative of the intensity signal and their intensity was quantified by integrating the area under the curve.

For western blotting, samples were transferred onto a PVDF membrane (Biorad) after SDS-PAGE Membranes were blocked in 5% (w/v) skim milk powder in Tris-buffered saline with 0.05 % (v/v) Tween-20 (TBS-T) before probing with primary antibodies diluted in blocking buffer overnight. Membranes were washed, incubated with secondary HRP-coupled antibodies for 1 h at room temperature and washed before probing with Amersham ECL Prime detection reagent (Cytiva) and chemiluminescence detection on a Fusion FX Edge (Vilber-Lourmat) or Amersham 600 (GE) imager. Densitometric quantification of western blots was performed with ImageJ Fiji using standard settings and statistical analyses were performed using GraphPad Prism version 10.

Antibodies used in this study: ATP13A1 (Proteintech, 16244-1-AP), EMC1 (Novus Biologicals, NBP2-59097 and NBP3-18427), EMC4 (Abcam, ab184544), EMC7 (Proteintech, 27550-1-AP), HA.11 (Biolegend, Poly9023), c-myc (Proteintech, 16286-1-AP), FLAG (Proteintech, 20543-1-AP), NWSHPQFEK 5A9F9 (Genscript, A01732), GFP (Proteintech, 50430-2-AP), Hsc70 (Santa Cruz Biotechnology, sc-7298), GAPDH-HRP (Proteintech, HRP-60004), anti-rabbit IgG-HRP (Santa Cruz Biotechnology, sc-2357), m-IgGκ BP-HRP, sc-516102).

## Acknowledgements

We thank Samuel Maiwald, Luca Stier, Dominik Magyar, Felix Johannsen, Emil Korzin, Josephine Botsch, Zebin Tong and Martin Haslbeck for support with experimental procedures; Leo Kiss and Filiz Civril Stocker for input on the manuscript; Daniel Bollschweiler and Tillman Schäfer for assistance with cryo-EM; Markus Oster for assistance with flow cytometry; and all members of the Schulman and the Feige labs for scholarly discussions. We thank Jonathan Weissman for consultation and valuable input on this project.

We would like to acknowledge the following research facilities at the MPI of Biochemistry: Cryo-EM Facility (RRID:SCR_025744), Mass Spectrometry Facility (RRID:SCR_025745), Imaging Facility (RRID:SCR_025739), as well as the TUM Pilot Plant for Industrial Biotechnology at the Technical University of Munich.

Work in MJF lab was funded by the DFG (FE 1581/5-1, to MJF and BB) and the European Union (ERC, DeCoDe, 101088970, to MJF).

Work in MS lab was supported by the European Union (ERC, OnTarget, 864068 to MS). MS is an incumbent of the Gilbert Omenn and Martha Darling Professorial Chair in Molecular Genetics.

Work in BAS lab was funded by the Max Planck Society, the European Union (ERC, UPSmeetMet, 101098161 to BAS), and the Deutsche Forschungsgemeinschaft (DFG, German Research Foundation) for Leibniz Prize SCHU 3196/1-1 to BAS.

Views and opinions expressed are however those of the author(s) only and do not necessarily reflect those of the European Union, the European Research Council or other funding agencies. Neither the European Union nor the granting authority can be held responsible for them.

EF was supported by a senior postdoctoral award from the Weizmann Institute of Science and is currently supported by the DFG (FE 2386/2-1) and the Center for Molecular Medicine Cologne (CMMC) Career Advancement Program (CAP-37).

CJK gratefully acknowledges Ph.D. fellowships from the Boehringer Ingelheim Fonds and the Studienstiftung des deutschen Volkes.

## Competing interests

BAS is a co-inventors of intellectual property related to DCUND1 inhibitors licensed to Cinsano. BAS is a member of the scientific advisory boards of Proxygen and Lyterian. The other authors declare no competing interests.

## Author contributions

Conceptualization: MJF, BB, BAS, CJK; protein purification: CJK; cryo-EM data collection and analysis: CJK, JRP; atomic model building: JRP; mass spectrometry: BS; yeast strain design and generation: EJF, CRL, SA, MS; yeast assays: CJK; mammalian cell line generation: CJK; mammalian co-IPs: IB, CJK; data analysis: CJK, BB, MS, MJF, BAS; funding acquisition: MJF, BAS, MS, BB; paper writing: CJK, MJF, BB, MS, BAS with input from all authors.

## Extended Data

**Extended Data Fig 1.**
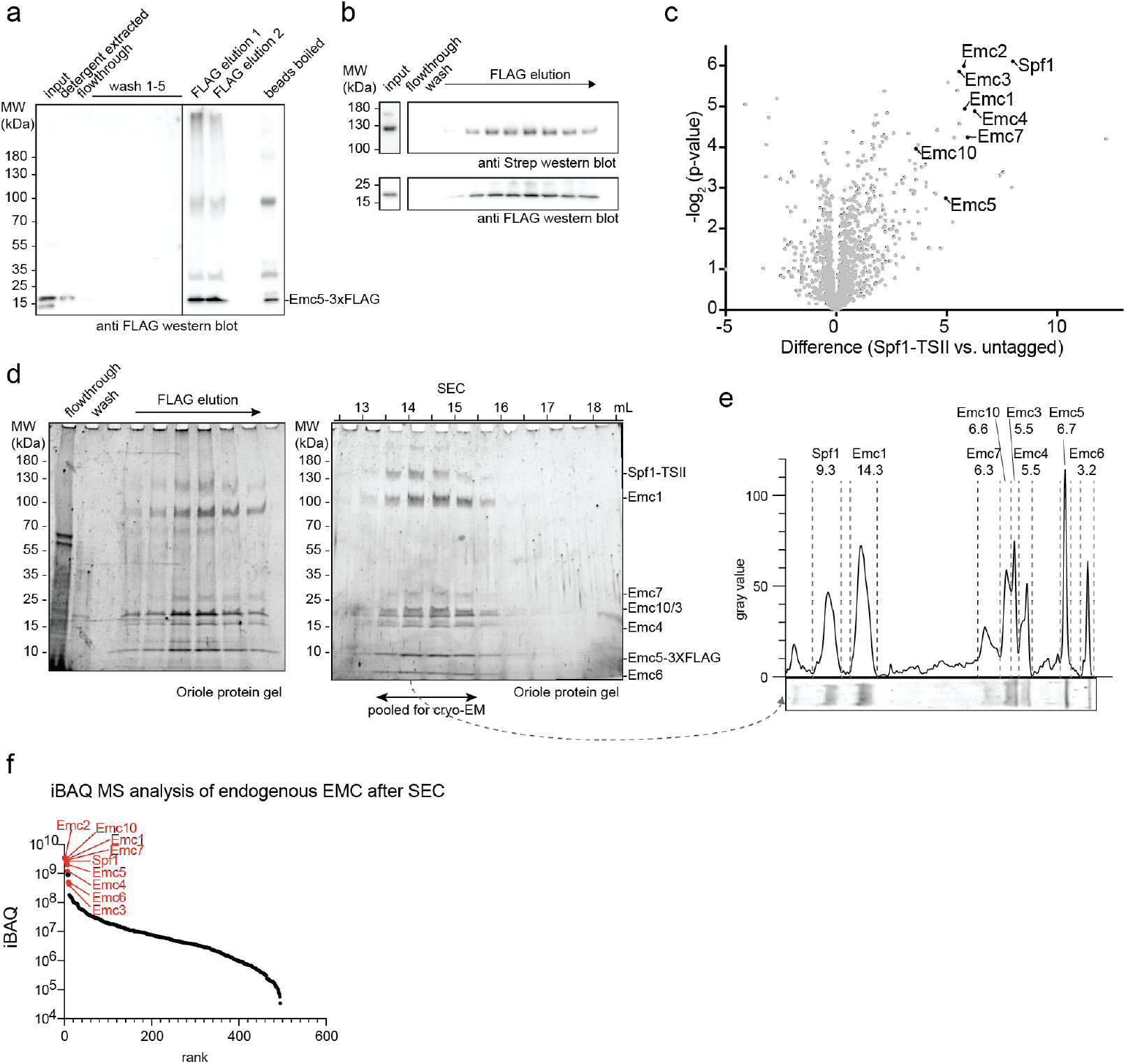
Validation of the EMC:Spf1 interaction and its stoichiometry at endogenous level in *S. cerevisiae*. **(A)** Anti FLAG western blot showing the different steps of the isolation procedure of Emc5-3XFLAG from *S. cerevisiae*. **(B)** Co-immunoprecipitation of endogenously tagged Spf1-TSII with Emc5-3XFLAG confirms the interaction at endogenous levels. **(C)** The interactome of endogenously TSII-tagged Spf1 validates the interaction with the EMC from the Spf1 side. **(D)** SDS-PAGE protein gels stained with Oriole stain of the large-scale purification of endogenous EMC from a 400 g yeast pellet by an anti-FLAG pulldown and size exclusion chromatography. The second gel is also shown in Fig. 1c and has been reproduced here for completeness. **(E)** Quantification of protein band intensities using densitometry of lane 4 of the SEC gel shown in d. The respective region was cropped and rotated by 90°, corrected for background signal and intensities were quantified. Individual peaks corresponding to Spf1 and the EMC subunits were marked and labeled with the integrated area under the curve. **(F)** iBAQ MS analysis of the endogenously isolated EMC validates Spf1 as most abundant interaction partner among the EMC subunits.

**Extended Data Fig 2.**
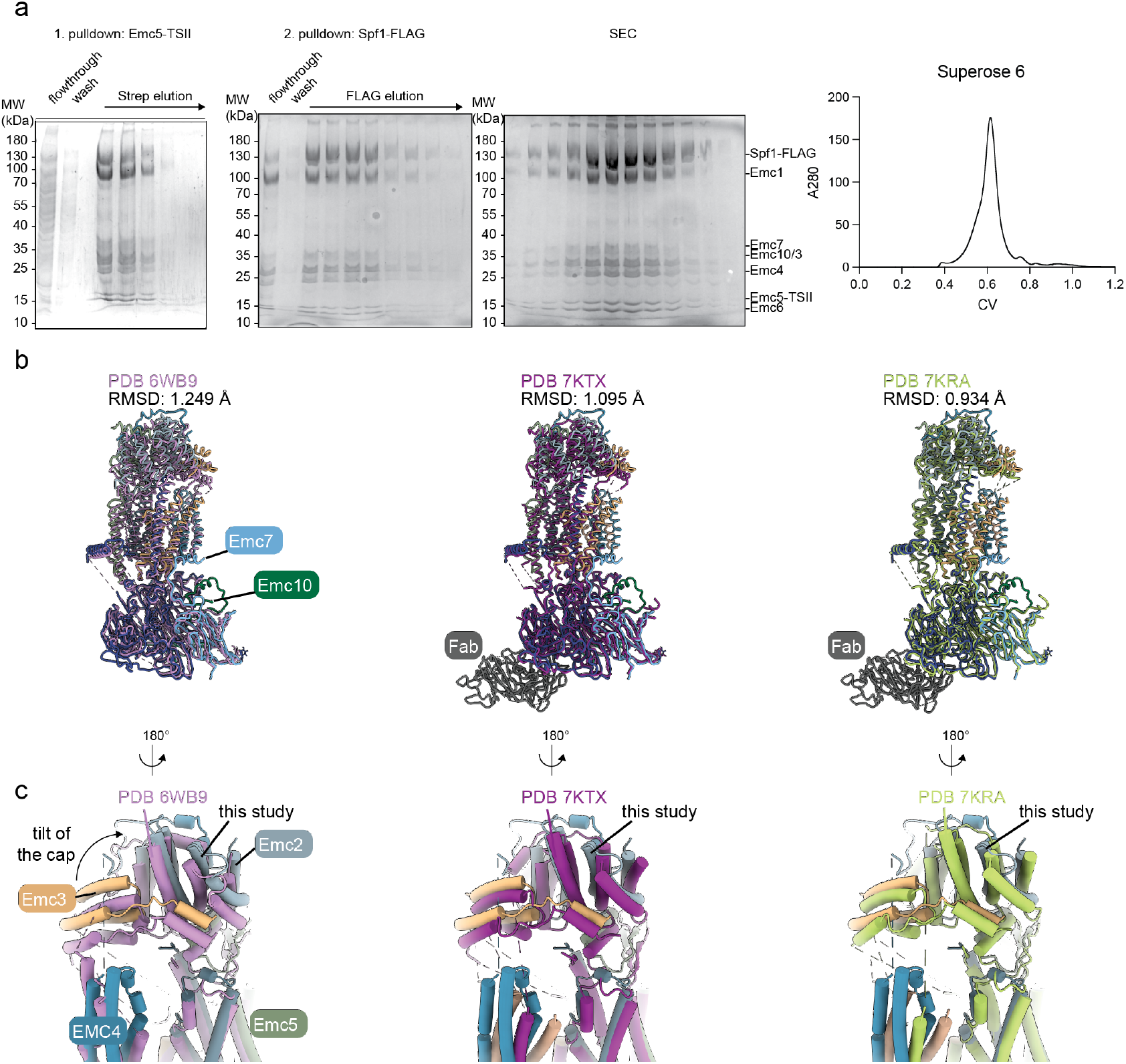
Purification of the recombinant EMC:Spf1 complex and comparison of the EMC to previously published structures. **(A)** Purification of the EMC:Spf1 complex from insect cells via a tandem pulldown, first on Emc5-TSII, second on Spf1-FLAG followed by size exclusion chromatography on a Superose 6 column. The chromatogram and a lane of the SEC gel are also shown in Fig. 2a and have been reproduced here for completeness. **(B)** Comparison of the EMC structure in complex with Spf1 to previously reported structures of the yeast EMC (PDBs: 6WB9, 7KTX, 7KRA). **(C)** View of the cytoplasmic cap of the EMC, which is shifted in the EMC:Spf1 complex in comparison the apo EMC. The arrow indicates the direction of the cap tilt.

**Extended Data Fig 3.**
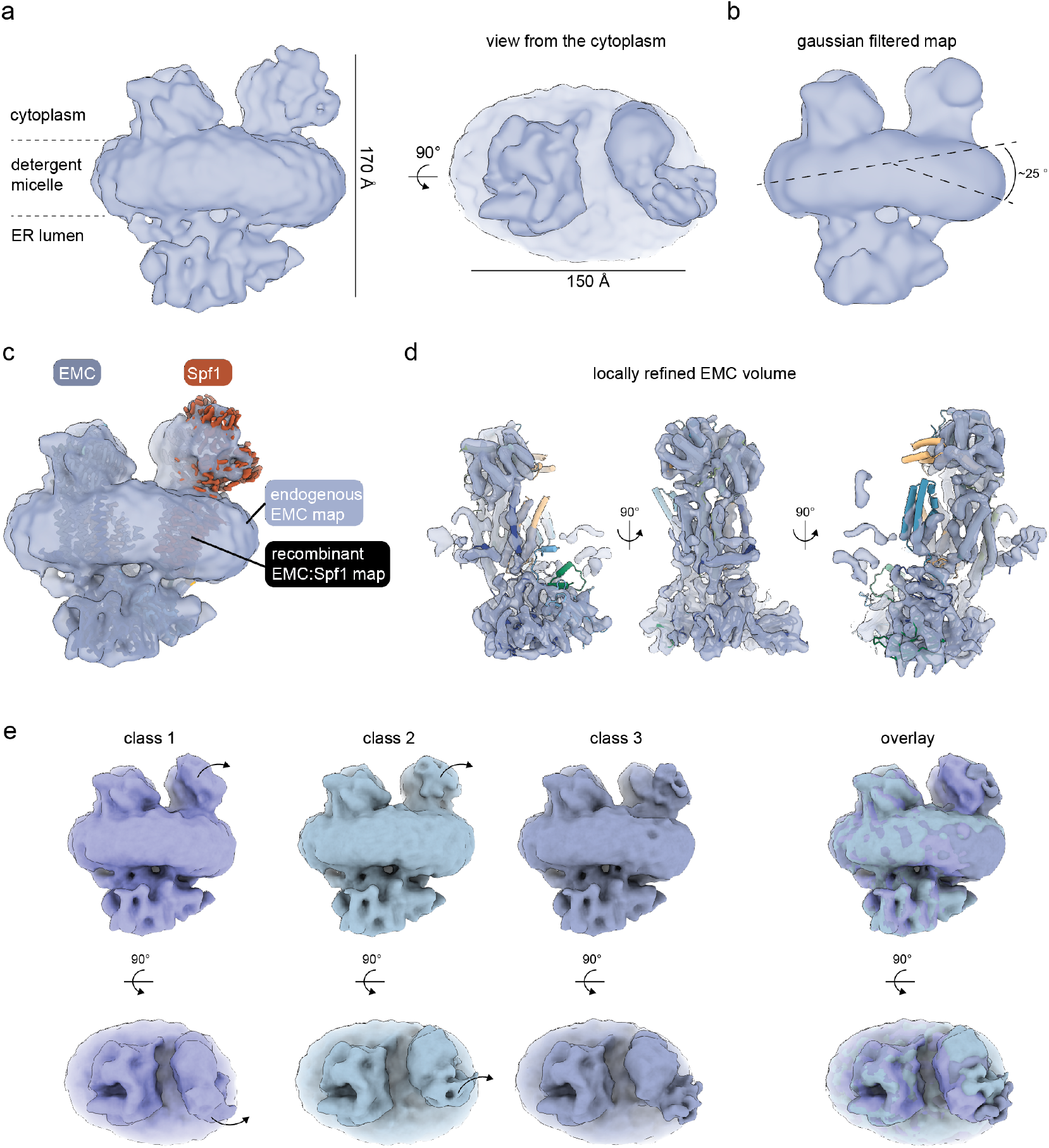
Cryo-EM map of the endogenous EMC:Spf1 from yeast. **(A)** Reconstruction of the endogenous EMC:Spf1 volume after 3D flexible refinement, showing an EMC-like density (left) and an additional density (right) that likely corresponds to Spf1. **(B)** The endogenous EMC:Spf1 complex introduces a bend in the detergent micelle to a similar extend as the recombinant complex. **(C)** Overlay of the maps of the EMC obtained from endogenous sources (grey) and the side-chain resolution map of the recombinantly expressed EMC:Spf1 supercomplex (color coded according to subunit identity) showing that the recombinant structure is fully consistent with the endogenous map. **(D)** Local refinement of the EMC component of the endogenous map shows secondary structure elements and consistency with the structural model that was derived from the recombinant EMC:Spf1 complex and is docked into the density here. **(E)** The endogenous EMC:Spf1 complex shows conformational heterogeneity in the Spf1 component, particularly density corresponding to the N and P-domains displays substantial flexibility. Three classes of the final consensus set of particles are shown and differences between the classes are indicated by arrows. An overlay of all classes is shown on the right.

**Extended Data Fig 4.**
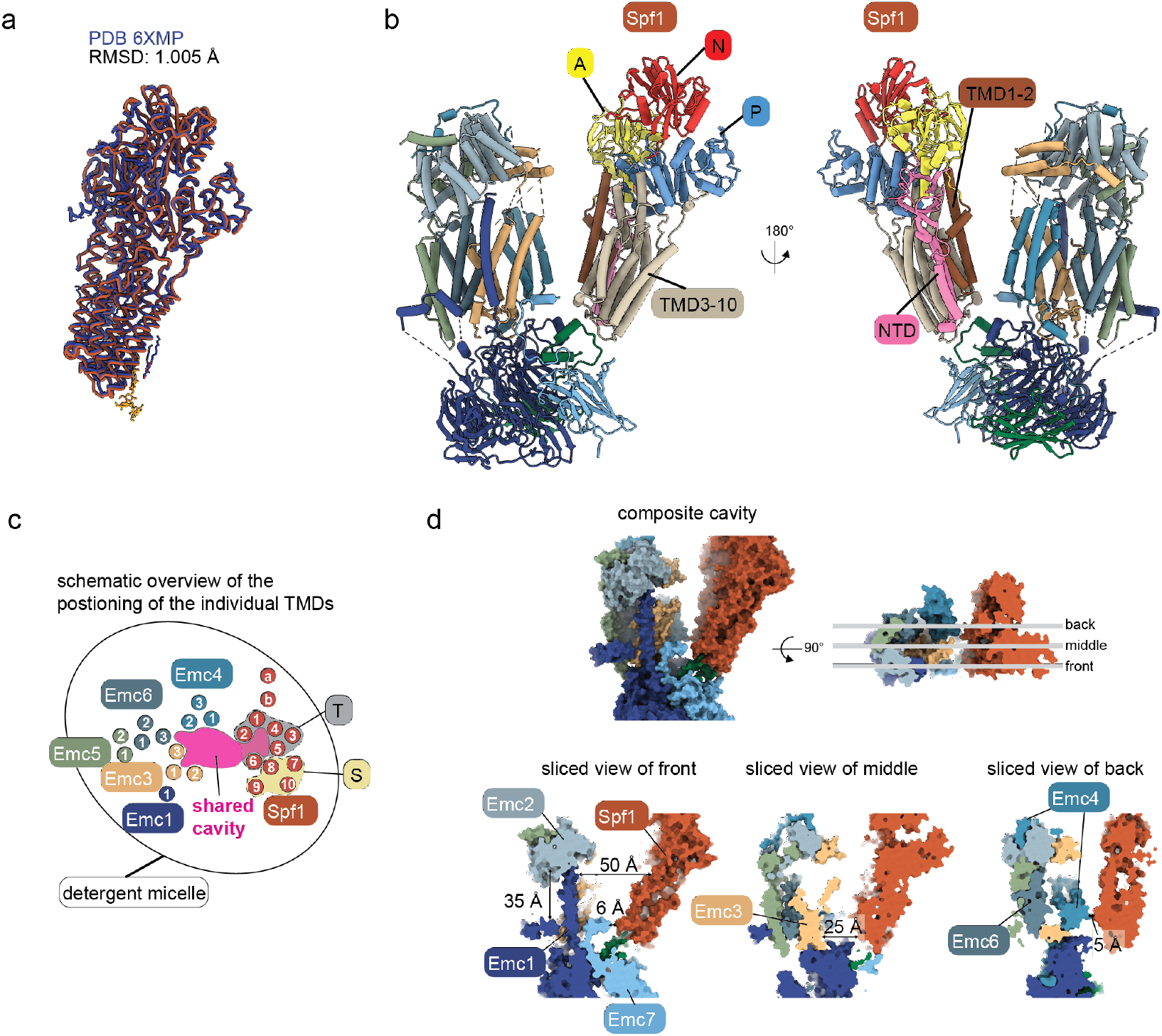
Domain architecture of Spf1 in complex with the EMC and details of the composite intramembrane cavity. **(A)** Comparison of the Spf1 structure in complex with EMC to a previously reported structure of apo Spf1 (PDB 6XMP). **(B)** Model of the EMC:Spf1 complex color coded according to Spf1 domain identity, featuring the actuator (A), phosphorylation (P), nucleotide binding (N) domains as well as the N-terminal domain (NTD). The substrate binding cleft is located between transmembrane domains colored in brown (TMD1-2) and in sand (TMD3-10). Arrangement of the transport (T; TMD1-6) and support domain (S; TMD7-10) is outlined in panel c. **(C)** Schematic overview of the arrangement of the TMDs of the EMC and Spf1 in the the detergent micelle and highlighting the area of the composite intramembrane cavity enclosed by these two protein complexes and the architecture of Spf1’s TMDs arranged into the transport (T) and support (S) domains. **(D)** Detailed views of the composite cavity enclosed by the EMC and Spf1. The model of the EMC:Spf1 complex is shown in surface representation from the membrane and the top. In the top view the planes are indicated where the sections (right) were taken.

**Extended Data Fig 5.**
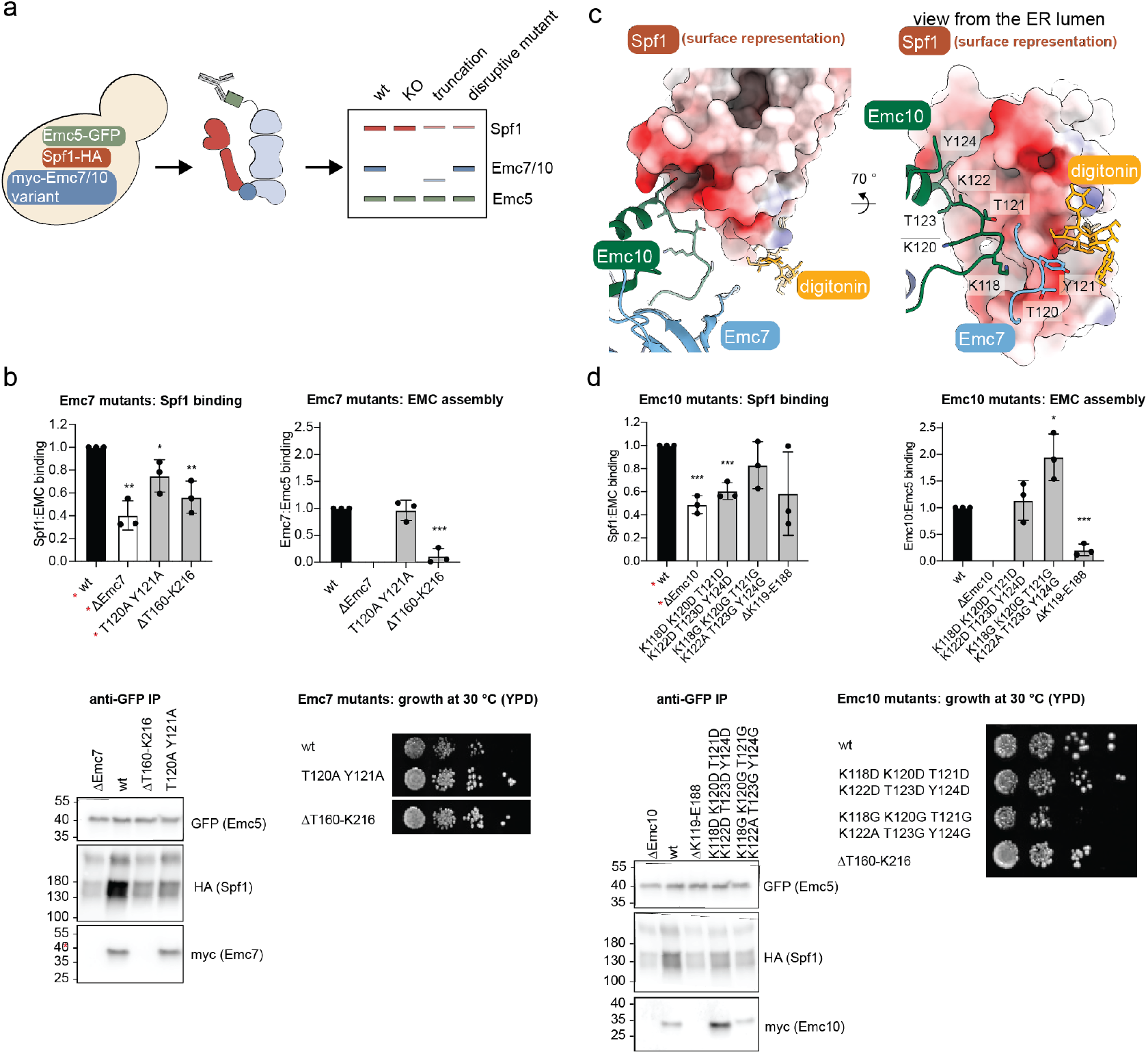
Validating the lumenal dock by mutagenesis of Emc7 and Emc10 in an endogenous setup in *S. cerevisiae*. **(A)** Schematic depiction of the experimental setup to validate the lumenal interface in an endogenous yeast setting. Emc5 was tagged C-terminally with a GFP, Spf1 with a triple HA-tag. Emc7 or Emc10 were either deleted or replaced at their endogenous locus with a version containing an N-terminal myc-tag, located after the signal sequence. The GFP was used as an affinity handle for co-IP experiments and binding of Spf1 and the Emc7/10 variant was assessed by western blot against the HA or myc tag, respectively. **(B)** Overview of the effects of all mutations in Emc7 on the Emc-Spf1 interaction and overall EMC assembly. Binding was assessed in at least three independent replicates and normalized to the respective wt. Binding was assessed in three independent replicates and normalized to the respective wt. (mean ± standard deviation, unpaired Student’s t-test * p < 0.05, ** p < 0.01, *** p < 0.005, **** p < 0.0001). A part of this data is also shown in Fig. 3c (indicated with red asterisks). Western blots of one representative replicate of anti-GFP IP are shown. A growth assay was performed to exclude effects of the Emc7 variants on yeast growth under standard conditions (YPD, 30 °C). **(C)** View of the lumenal dock with Spf1 depicted as surface rendering, colored according to electrostatic potential. The view from the ER lumenal side shows the negatively charged lumenal surface of Spf1 that is contacted by Emc10 (green). Residues that were mutated in Emc10 are labeled. **(D)** Overview of the effects of all mutations in Emc10 on the Emc-Spf1 interaction and overall EMC assembly as in panel b.

**Extended Data Fig 6.**
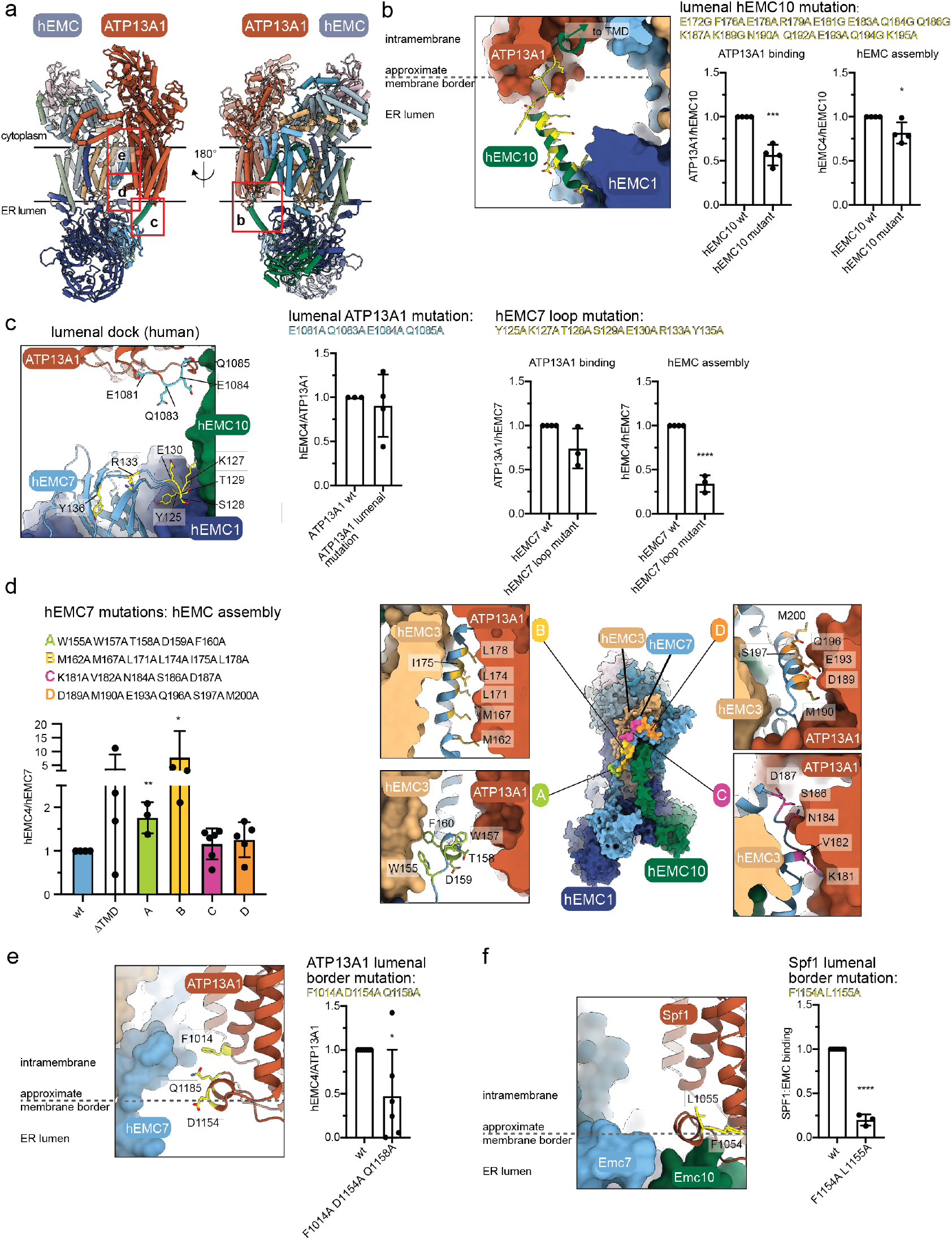
Mutational dissection of the Alphafold3 model of the hEMC:ATP13A1 complex. **(A)** Predicted model of the hEMC:ATP13A1 complex from two views as shown in Fig. 4b. To orient the reader, regions that were mutated and are shown in detail in the following panels are marked with red frames and the letters labeling that panel are included. Approximate membrane borders are marked with black lines. **(B)** Mutation of the lumenal linker and helix region in hEMC10 affects ATP13A1 binding in cells and mildly compromises hEMC assembly. Mutated residues are highlighted in yellow. Binding was assessed via co-IP on hEMC10 in four independent replicates and normalized to the wt. (mean ± standard deviation, unpaired Student’s t-test * p < 0.05, ** p < 0.01, *** p < 0.005, **** p < 0.0001). **(C)** View of the lumenal dock in the human complex. Aquamarine: Acidic residues in ATP13A1 that were mutated and had no effect on hEMC binding (bar graph on the right). Yellow: Residues in the region of hEMC7, corresponding to the Emc7 loop in yeast, whose mutation resulted in reduced association with the hEMC yet had no significant effect on hEMC binding (bar graphs on the bottom). Binding was assessed via co-IP on ATP13A1 or hEMC7, respectively in three (hEMC7 mutation) or four (ATP13A1 mutation) independent replicates and normalized to the respective wt. (mean ± standard deviation, unpaired Student’s t-test * p < 0.05, ** p < 0.01, *** p < 0.005, **** p < 0.0001). **(D)** Left: Effect of mutations in hEMC7 shown in Fig. 4c on hEMC assembly. Deletion of the TMD or mutation of subregions thereof had a stabilizing or no effect. Binding was assessed via co-IP on hEMC7 in at least three independent replicates (n=3 for A, B; n=4 for βTMD, n=5 for D, n=6 for C) and normalized to the wt. (mean ± standard deviation, unpaired Student’s t-test * p < 0.05, ** p < 0.01, *** p < 0.005, **** p < 0.0001). Right: detailed depiction of the residues mutated in the individual segments of hEMC7. **(E)** Mutation of selected residues in ATP13A1 (highlighted in yellow) at the border between the transmembrane region and ER lumen destabilize complex formation with the hEMC. Binding was assessed via co-IP on ATP13A1 in six independent replicates and normalized to the wt. (mean ± standard deviation, unpaired Student’s t-test * p < 0.05, ** p < 0.01, *** p < 0.005, **** p < 0.0001). **(F)** Mutation of residues in Spf1 at the border between the transmembrane region and ER lumen, corresponding to the mutation shown in e in ATP13A1. Binding was assessed via co-IP on Spf1 expressed recombinantly in insect cells in three independent replicates and normalized to the wt. (mean ± standard deviation, unpaired Student’s t-test * p < 0.05, ** p < 0.01, *** p < 0.005, **** p < 0.0001).

**Extended Data Fig 7.**
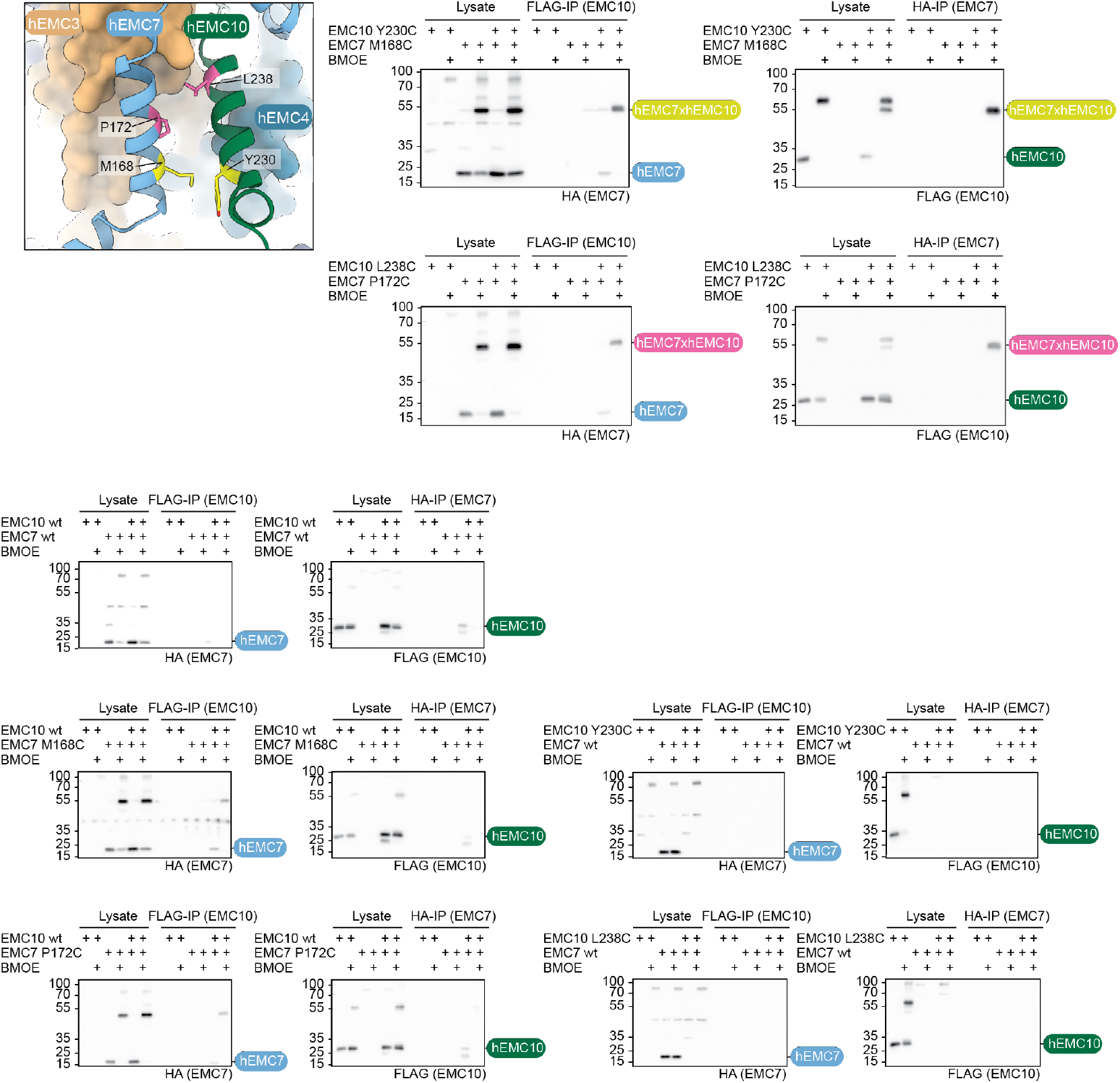
Intramembrane cysteine crosslinking between hEMC7 and hEMC10 confirms the predicted TMD interface. Single residues in hEMC7 (M168, P172) and hEMC10 (Y230, L238) were replaced with cysteines and in-cell crosslinking was performed using BMOE. Crosslinked adducts were isolated by IP and analyzed on a western blot, confirming the presence of a successful cysteine crosslink between the two tagged subunits. hEMC10 contains 4 cysteines in its wt sequence. To exclude the possibility of bands arising from crosslinking with other cysteines than the ones introduced or unspecific interaction, a combination of hEMC7 and hEMC10 without any further Cys mutation was assessed and also each Cys-variant was tested with the wt of the other protein.

**Extended Data Fig 8.**
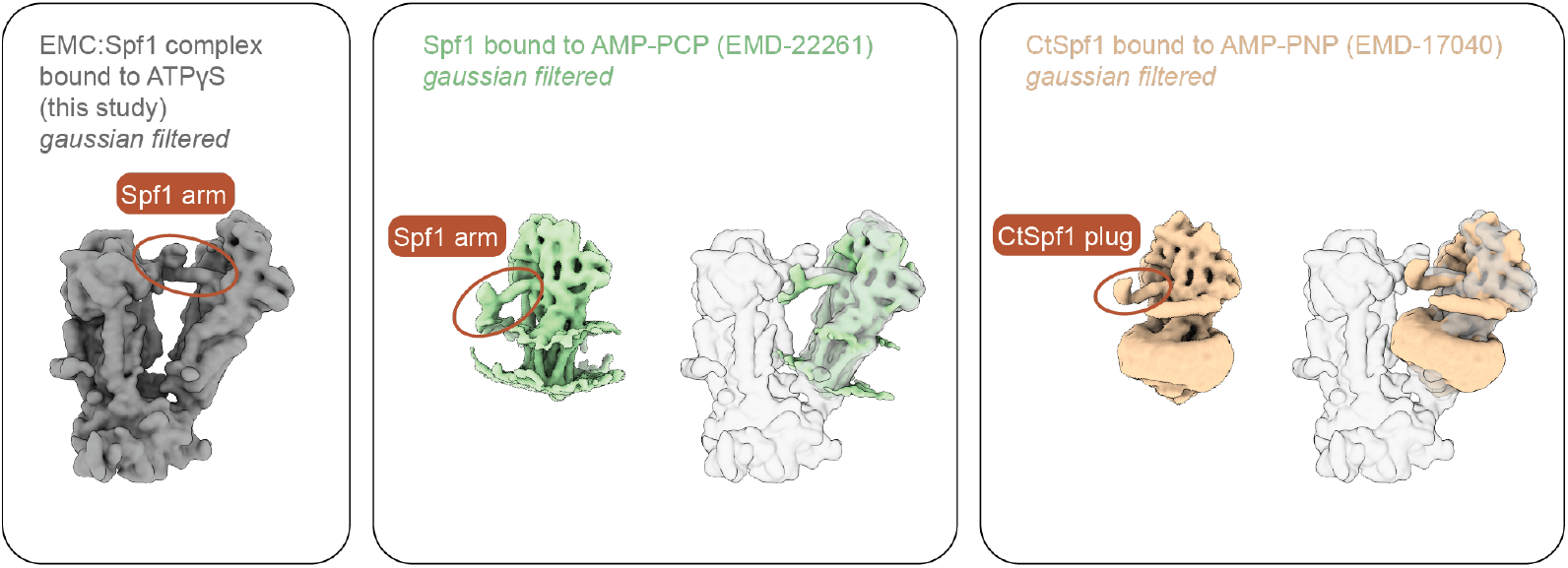
Consistent positioning of the arm domain of Spf1 in the E1-ATP state. The Spf1 arm domain is stabilized in the E1-ATP state in the EMC:Spf1 complex, allowing structural characterization and modeling. A comparison to lower resolution reports of the Spf1 arm^26^ or equivalent CtSpf1 “plug” domain^31^ shows consistent positioning in the respective E1-ATP state. A gaussian filter was applied to all maps to allow comparison at a similar resolution level.

**Extended Data Fig 9.**
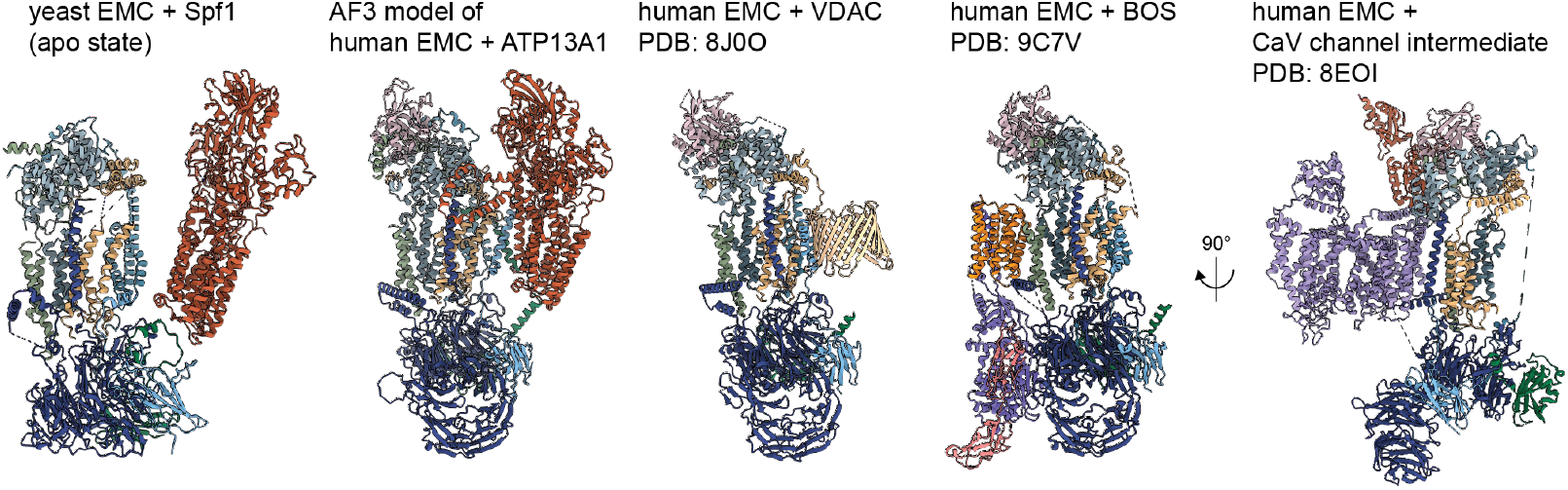
Comparison of the EMC:Spf1 complex from yeast and the predicted hEMC:ATP13A1 complex to other EMC complexes that have been structurally characterized^**13,18,55**^.

## Notes

### Competing Interest Statement

BAS is a co-inventor of intellectual property related to DCUND1 inhibitors licensed to Cinsano. BAS is a member of the scientific advisory boards of Proxygen and Lyterian. The other authors declare no competing interests.

### Summary of Updates

The author affiliations were updated and formatting was adjusted.

## References

1 Pleiner, T. et al. A selectivity filter in the ER membrane protein complex limits protein misinsertion at the ER. J Cell Biol 222 (2023). 10.1083/jcb.202212007

2 Hegde, R. S. & Keenan, R. J. The mechanisms of integral membrane protein biogenesis. Nat Rev Mol Cell Biol 23, 107–124 (2022). 10.1038/s41580-021-00413-2

3 Weill, U., Cohen, N., Fadel, A., Ben-Dor, S. & Schuldiner, M. Protein Topology Prediction Algorithms Systematically Investigated in the Yeast Saccharomyces cerevisiae. Bioessays 41, e1800252 (2019). 10.1002/bies.201800252

4 Walter, P., Gilmore, R. & Blobel, G. Protein translocation across the endoplasmic reticulum. Cell 38, 5–8 (1984). https://doi.org/0092-8674(84)90520-8 [pii]

5 Schuldiner, M. et al. The GET complex mediates insertion of tail-anchored proteins into the ER membrane. Cell 134, 634–645 (2008). 10.1016/j.cell.2008.06.025

6 Aviram, N. et al. The SND proteins constitute an alternative targeting route to the endoplasmic reticulum. Nature 540, 134–138 (2016). 10.1038/nature20169

7 Stirling, C. J., Rothblatt, J., Hosobuchi, M., Deshaies, R. & Schekman, R. Protein translocation mutants defective in the insertion of integral membrane proteins into the endoplasmic reticulum. Mol Biol Cell 3, 129–142 (1992). 10.1091/mbc.3.2.129

8 Green, N., Fang, H. & Walter, P. Mutants in three novel complementation groups inhibit membrane protein insertion into and soluble protein translocation across the endoplasmic reticulum membrane of Saccharomyces cerevisiae. J Cell Biol 116, 597–604 (1992). 10.1083/jcb.116.3.597

9 Oliver, J., Jungnickel, B., Görlich, D., Rapoport, T. & High, S. The Sec61 complex is essential for the insertion of proteins into the membrane of the endoplasmic reticulum. FEBS Lett 362, 126–130 (1995). 10.1016/0014-5793(95)00223-v

10 McGilvray, P. T. et al. An ER translocon for multi-pass membrane protein biogenesis. eLife 9 (2020). 10.7554/eLife.56889

11 Page, K. R. et al. Role of a holo-insertase complex in the biogenesis of biophysically diverse ER membrane proteins. Mol Cell (2024). 10.1016/j.molcel.2024.08.005

12 Jonikas, M. C. et al. Comprehensive characterization of genes required for protein folding in the endoplasmic reticulum. Science 323, 1693–1697 (2009). 10.1126/science.1167983

13 Page, K. R. et al. Role of a holo-insertase complex in the biogenesis of biophysically diverse ER membrane proteins. Mol Cell 84, 3302–3319 e3311 (2024). 10.1016/j.molcel.2024.08.005

14 Chitwood, P. J., Juszkiewicz, S., Guna, A., Shao, S. & Hegde, R. S. EMC Is Required to Initiate Accurate Membrane Protein Topogenesis. Cell (2018). 10.1016/j.cell.2018.10.009

15 Guna, A., Volkmar, N., Christianson, J. C. & Hegde, R. S. The ER membrane protein complex is a transmembrane domain insertase. Science 359, 470–473 (2018). 10.1126/science.aao3099

16 Hegde, R. S. The Function, Structure, and Origins of the ER Membrane Protein Complex. Annu Rev Biochem 91, 651–678 (2022). 10.1146/annurev-biochem-032620-104553

17 Shurtleff, M. J. et al. The ER membrane protein complex interacts cotranslationally to enable biogenesis of multipass membrane proteins. Elife 7 (2018). 10.7554/eLife.37018

18 Chen, Z. et al. EMC chaperone-Ca(V) structure reveals an ion channel assembly intermediate. Nature 619, 410–419 (2023). 10.1038/s41586-023-06175-5

19 Klose, C. J. et al. The EMC acts as a chaperone for membrane proteins. Nature Communications 16, 7097 (2025). 10.1038/s41467-025-62109-x

20 Christianson, J. C. et al. Defining human ERAD networks through an integrative mapping strategy. Nat Cell Biol 14, 93–105 (2011). 10.1038/ncb2383

21 Okreglak, V. & Walter, P. The conserved AAA-ATPase Msp1 confers organelle specificity to tail-anchored proteins. Proc Natl Acad Sci U S A 111, 8019–8024 (2014). 10.1073/pnas.1405755111

22 Chen, Y. C. et al. Msp1/ATAD1 maintains mitochondrial function by facilitating the degradation of mislocalized tail-anchored proteins. EMBO J 33, 1548–1564 (2014). 10.15252/embj.201487943

23 Delgado, J. M., Shepard, L. W., Lamson, S. W., Liu, S. L. & Shoemaker, C. J. The ER membrane protein complex restricts mitophagy by controlling BNIP3 turnover. EMBO J 43, 32–60 (2024). 10.1038/s44318-023-00006-z

24 Gungor, B., Flohr, T., Garg, S. G. & Herrmann, J. M. The ER membrane complex (EMC) can functionally replace the Oxa1 insertase in mitochondria. PLoS Biol 20, e3001380 (2022). 10.1371/journal.pbio.3001380

25 McKenna, M. J., Adams, B. M., Chu, V., Paulo, J. A. & Shao, S. ATP13A1 prevents ERAD of folding-competent mislocalized and misoriented proteins. Mol Cell 82, 4277–4289 e4210 (2022). 10.1016/j.molcel.2022.09.035

26 McKenna, M. J. et al. The endoplasmic reticulum P5A-ATPase is a transmembrane helix dislocase. Science 369 (2020). 10.1126/science.abc5809

27 Ji, J. et al. An ATP13A1-assisted topogenesis pathway for folding multi-spanning membrane proteins. Mol Cell 84, 1917-1931.e1915 (2024). 10.1016/j.molcel.2024.04.010

28 McKenna, M. J., Adams, B. M., Chu, V., Paulo, J. A. & Shao, S. ATP13A1 prevents ERAD of folding-competent mislocalized and misoriented proteins. Mol Cell 82, 4277-4289.e4210 (2022). 10.1016/j.molcel.2022.09.035

29 Post, R. L., Hegyvary, C. & Kume, S. Activation by adenosine triphosphate in the phosphorylation kinetics of sodium and potassium ion transport adenosine triphosphatase. J Biol Chem 247, 6530–6540 (1972).

30 Albers, R. W. Biochemical aspects of active transport. Annu Rev Biochem 36, 727–756 (1967). 10.1146/annurev.bi.36.070167.003455

31 Li, P. et al. The structure and function of P5A-ATPases. Nat Commun 15, 9605 (2024). 10.1038/s41467-024-53757-6

32 Volkmar, N. et al. The ER membrane protein complex promotes biogenesis of sterol-related enzymes maintaining cholesterol homeostasis. J Cell Sci 132 (2019). 10.1242/jcs.223453

33 Krumpe, K. et al. Ergosterol content specifies targeting of tail-anchored proteins to mitochondrial outer membranes. Mol Biol Cell 23, 3927–3935 (2012). 10.1091/mbc.E11-12-0994

34 Sorensen, D. M. et al. The P5A ATPase Spf1p is stimulated by phosphatidylinositol 4-phosphate and influences cellular sterol homeostasis. Mol Biol Cell 30, 1069–1084 (2019). 10.1091/mbc.E18-06-0365

35 Ji, J. et al. An ATP13A1-assisted topogenesis pathway for folding multi-spanning membrane proteins. Mol Cell 84, 1917–1931 e1915 (2024). 10.1016/j.molcel.2024.04.010

36 Hegde, R. S. Getting membrane proteins into shape. Mol Cell 84, 1821–1823 (2024). 10.1016/j.molcel.2024.04.024

37 Bai, L., You, Q., Feng, X., Kovach, A. & Li, H. Structure of the ER membrane complex, a transmembrane-domain insertase. Nature 584, 475–478 (2020). 10.1038/s41586-020-2389-3

38 Miller-Vedam, L. E. et al. Structural and mechanistic basis of the EMC-dependent biogenesis of distinct transmembrane clients. Elife 9 (2020). 10.7554/eLife.62611

39 O’Donnell, J. P. et al. The architecture of EMC reveals a path for membrane protein insertion. Elife 9 (2020). 10.7554/eLife.57887

40 Anghel, S. A., McGilvray, P. T., Hegde, R. S. & Keenan, R. J. Identification of Oxa1 Homologs Operating in the Eukaryotic Endoplasmic Reticulum. Cell Rep 21, 3708–3716 (2017). 10.1016/j.celrep.2017.12.006

41 Wu, H., Smalinskaite, L. & Hegde, R. S. EMC rectifies the topology of multipass membrane proteins. Nat Struct Mol Biol 31, 32–41 (2024). 10.1038/s41594-023-01120-6

42 Hegde, R. S. & Keenan, R. J. A unifying model for membrane protein biogenesis. Nat Struct Mol Biol 31, 1009–1017 (2024). 10.1038/s41594-024-01296-5

43 Pleiner, T. et al. Structural basis for membrane insertion by the human ER membrane protein complex. Science 369, 433–436 (2020). 10.1126/science.abb5008

44 Wu, H. & Hegde, R. S. Mechanism of signal-anchor triage during early steps of membrane protein insertion. Mol Cell 83, 961–973 e967 (2023). 10.1016/j.molcel.2023.01.018

45 Diamantopoulou, A. et al. Loss-of-function mutation in Mirta22/Emc10 rescues specific schizophrenia-related phenotypes in a mouse model of the 22q11.2 deletion. Proc Natl Acad Sci U S A 114, E6127–E6136 (2017). 10.1073/pnas.1615719114

46 Zhou, Y. et al. EMC10 governs male fertility via maintaining sperm ion balance. J Mol Cell Biol 10, 503–514 (2018). 10.1093/jmcb/mjy024

47 Umair, M. et al. EMC10 homozygous variant identified in a family with global developmental delay, mild intellectual disability, and speech delay. Clin Genet 98, 555–561 (2020). 10.1111/cge.13842

48 Shao, D. D. et al. A recurrent, homozygous EMC10 frameshift variant is associated with a syndrome of developmental delay with variable seizures and dysmorphic features. Genet Med 23, 1158–1162 (2021). 10.1038/s41436-021-01097-x

49 Kaiyrzhanov, R. et al. Biallelic loss of EMC10 leads to mild to severe intellectual disability. Ann Clin Transl Neurol 9, 1080–1089 (2022). 10.1002/acn3.51602

50 Chen, K. et al. EMC10 modulates hepatic ER stress and steatosis in an isoform-specific manner. J Hepatol 81, 479–491 (2024). 10.1016/j.jhep.2024.03.047

51 Abramson, J. et al. Accurate structure prediction of biomolecular interactions with AlphaFold 3. Nature 630, 493–500 (2024). 10.1038/s41586-024-07487-w

52 Qin, Q., Zhao, T., Zou, W., Shen, K. & Wang, X. An Endoplasmic Reticulum ATPase Safeguards Endoplasmic Reticulum Identity by Removing Ectopically Localized Mitochondrial Proteins. Cell Rep 33, 108363 (2020). 10.1016/j.celrep.2020.108363

53 Tipper, D. J. & Harley, C. A. Yeast genes controlling responses to topogenic signals in a model transmembrane protein. Mol Biol Cell 13, 1158–1174 (2002). 10.1091/mbc.01-10-0488

54 Yang, X. et al. ATP13A1 engages SEC61 to facilitate substrate-specific translocation. Sci Adv 11, eadt1346 (2025). 10.1126/sciadv.adt1346

55 Li, M. et al. Structural insights into human EMC and its interaction with VDAC. Aging (Albany NY) 16, 5501–5525 (2024). 10.18632/aging.205660

56 Weissmann, F. et al. biGBac enables rapid gene assembly for the expression of large multisubunit protein complexes. Proc Natl Acad Sci U S A 113, E2564–2569 (2016). 10.1073/pnas.1604935113

57 Knop, M. et al. Epitope tagging of yeast genes using a PCR-based strategy: more tags and improved practical routines. Yeast 15, 963–972 (1999). 10.1002/(SICI)1097-0061(199907)15:10B<963::AID-YEA399>3.0.CO;2-W

58 Janke, C. et al. A versatile toolbox for PCR-based tagging of yeast genes: new fluorescent proteins, more markers and promoter substitution cassettes. Yeast 21, 947–962 (2004). 10.1002/yea.1142

59 Gietz, R. D. & Woods, R. A. Transformation of yeast by lithium acetate/single-stranded carrier DNA/polyethylene glycol method. Methods Enzymol 350, 87–96 (2002). 10.1016/s0076-6879(02)50957-5

60 Gorlich, D. & Rapoport, T. A. Protein translocation into proteoliposomes reconstituted from purified components of the endoplasmic reticulum membrane. Cell 75, 615–630 (1993). 10.1016/0092-8674(93)90483-7

61 Cox, J. & Mann, M. MaxQuant enables high peptide identification rates, individualized p.p.b.-range mass accuracies and proteome-wide protein quantification. Nat Biotechnol 26, 1367–1372 (2008). 10.1038/nbt.1511

62 Tyanova, S. et al. The Perseus computational platform for comprehensive analysis of (prote)omics data. Nat Methods 13, 731–740 (2016). 10.1038/nmeth.3901

63 Mastronarde, D. N. Automated electron microscope tomography using robust prediction of specimen movements. J Struct Biol 152, 36–51 (2005). 10.1016/j.jsb.2005.07.007

64 Scheres, S. H. A Bayesian view on cryo-EM structure determination. J Mol Biol 415, 406–418 (2012). 10.1016/j.jmb.2011.11.010

65 Scheres, S. H. RELION: implementation of a Bayesian approach to cryo-EM structure determination. J Struct Biol 180, 519–530 (2012). 10.1016/j.jsb.2012.09.006

66 Rohou, A. & Grigorieff, N. CTFFIND4: Fast and accurate defocus estimation from electron micrographs. J Struct Biol 192, 216–221 (2015). 10.1016/j.jsb.2015.08.008

67 Zhang, K. Gautomatch.

68 Sanchez-Garcia, R. et al. DeepEMhancer: a deep learning solution for cryo-EM volume post-processing. Commun Biol 4, 874 (2021). 10.1038/s42003-021-02399-1

69 Liebschner, D. et al. Macromolecular structure determination using X-rays, neutrons and electrons: recent developments in Phenix. Acta Crystallogr D Struct Biol 75, 861–877 (2019). 10.1107/S2059798319011471

70 Punjani, A., Rubinstein, J. L., Fleet, D. J. & Brubaker, M. A. cryoSPARC: algorithms for rapid unsupervised cryo-EM structure determination. Nat Methods 14, 290–296 (2017). 10.1038/nmeth.4169

71 Punjani, A. & Fleet, D. J. 3DFlex: determining structure and motion of flexible proteins from cryo-EM. Nat Methods 20, 860–870 (2023). 10.1038/s41592-023-01853-8

72 Jumper, J. et al. Highly accurate protein structure prediction with AlphaFold. Nature 596, 583–589 (2021). 10.1038/s41586-021-03819-2

73 Varadi, M. et al. AlphaFold Protein Structure Database in 2024: providing structure coverage for over 214 million protein sequences. Nucleic Acids Res 52, D368–D375 (2024). 10.1093/nar/gkad1011

74 Pettersen, E. F. et al. UCSF Chimera--a visualization system for exploratory research and analysis. J Comput Chem 25, 1605–1612 (2004). 10.1002/jcc.20084

75 Adams, P. D. et al. PHENIX: a comprehensive Python-based system for macromolecular structure solution. Acta Crystallogr D Biol Crystallogr 66, 213–221 (2010). 10.1107/S0907444909052925

76 Emsley, P., Lohkamp, B., Scott, W. G. & Cowtan, K. Features and development of Coot. Acta Crystallogr D Biol Crystallogr 66, 486–501 (2010). 10.1107/S0907444910007493

77 Goddard, T. D. et al. UCSF ChimeraX: Meeting modern challenges in visualization and analysis. Protein Sci 27, 14–25 (2018). 10.1002/pro.3235

78 Ran, F. A. et al. Genome engineering using the CRISPR-Cas9 system. Nat Protoc 8, 2281–2308 (2013). 10.1038/nprot.2013.143

79 Schindelin, J. et al. Fiji: an open-source platform for biological-image analysis. Nat Methods 9, 676–682 (2012). 10.1038/nmeth.2019

